# Epigenetic regulation of Leukocyte associated immunoglobulin-like receptors 1 and 2 by interferon signaling in macrophages and T cells

**DOI:** 10.1101/2023.07.31.551385

**Authors:** Hannah K. Dorando, Evan C. Mutic, Joanna Y. Li, Ezri P. Perrin, Mellisa Wurtz, Chaz C. Quinn, Jacqueline E. Payton

**Author notes:** Contributed equally to authorship (Hannah K. Dorando, Evan C. Mutic, Joanna Y. Li). Correspondence: Jacqueline E. Payton 660 S. Euclid Ave, Box 8118 St. Louis, MO 63110.

## Abstract

**Background:** Inhibitory immune receptors are important for maintaining immune homeostasis. We recently identified epigenetic alterations in two members of this group, LAIR1 and LAIR2, in patients with inflammatory tissue damage and recurrent skin and soft tissue infections. We therefore hypothesized that the expression of LAIR1 and LAIR2 may be controlled by immune stimuli acting on discrete transcriptional regulatory elements.

**Methods:** We used flow cytometry, qRT-PCR, and RNAseq to assay LAIR1 and LAIR2 expression in human and murine immune cell subsets at baseline and post-treatment with immune mediators, including type I and II interferons, tumor necrosis factor-alpha (TNF-ɑ), and lipopolysaccharide (LPS). Using chromatin immunoprecipitation sequencing (ChIP-seq), we identified candidate transcriptional regulatory elements of LAIR genes and evaluated their regulatory activity using luciferase reporters.

**Results:** Both human and murine macrophages significantly upregulate LAIR1 expression as they differentiate from monocytes to macrophages. In response to interferons, LAIR1 protein levels increase, while LPS causes a relative reduction. Regulatory elements flanking LAIR genes exhibit distinct patterns of enhancer activity with variable responses to immune stimuli. These responses are related to discrete sets of transcription factors in inflammatory pathways that correlate with cell-specific LAIR expression patterns. In addition, we identified *LAIR1* and *LAIR2* regulatory elements that act as foci of 3D genome interactions with other highly active regulatory elements.

**Conclusions:** Our findings define the complex regulatory landscapes of human and mouse LAIR genes and reveal new insights into the transcriptional regulatory mechanisms that control the expression of these important immune modulatory proteins.

## INTRODUCTION

Immune inhibitory receptors are critical regulators of immune cell function; they modulate immune response to infection while limiting autoimmunity and immunopathology. In this way, inhibitory receptors contribute to immune homeostasis and tolerance. Conversely, deregulation of these receptors leads to impaired immune response to infection, autoimmunity, and immunopathology^1–3^. Recently, this large group of receptors has garnered renewed interest as potential drug targets for treating autoimmune disease and cancer^4^. We previously identified a member of this group of proteins, leukocyte-associated immunoglobulin (Ig)-like receptors (LAIR2), as associated with advanced and histone deacetylase inhibitor-resistant cutaneous T-cell lymphoma (CTCL), a disease marked by inflammatory tissue damage and deficits in cellular immunity ^5^.

LAIR1 and LAIR2 are Ig-like inhibitory receptors that are expressed in immune cells. LAIR1 is membrane-bound and expressed in both human and mouse cells, while LAIR2 is secreted, lacks the inhibitory ITIM domain, and is expressed only by human cells. They have nearly identical Ig-like domains and bind to the same collagen-like domain-containing ligands ^6–8^. LAIR1’s two cytosolic ITIM domains transmit inhibitory signals via SHP1 and SHP2 protein phosphatases that negatively regulate cytokine production^9–13^. By binding to the same ligands, LAIR2 has been shown to block LAIR1 signaling, thereby acting as an inhibitor of the inhibitor ^6^.

The genes encoding LAIR proteins lie within the leukocyte receptor complex (LRC), on chromosome 19p13.4 in humans and chromosome 7qA1 in mice. In addition to LAIR genes, the LRC contains many others that encode Ig-like receptors, including killer cell Ig-like receptors (*KIR*) and leukocyte Ig-like receptors (*LILR*). Genetic comparisons across species indicate that an early duplication and inversion event involving the primitive LRC region included a progenitor LAIR gene as well as intergenic regions, which likely included gene regulatory elements. In the ensuing millennia, many additional duplication and deletion events occurred, resulting in differing sets of Ig-like genes across vertebrate phyla^14^. Despite these differences, including the absence of *LAIR2* in mice, *LAIR1* and *Lair1* have similar exon/intron organization and protein structure^15^. From a gene regulatory point of view, *LAIR1* and *LAIR2* present a fascinating case in which duplicated and highly homologous genes share a similar regulatory genomic neighborhood, yet their protein products act in opposition to each other. Such a situation suggests that distinct transcriptional control of each gene would be necessary to prevent co-expression and thus canceling each other out. Indeed, as we show here, transcription levels of *LAIR1* and *LAIR2* vary by cell type. LAIR1 is expressed in most immune cell subsets, but levels vary widely across subsets and differentiation states, in both human and mouse^10, 11, 16^. LAIR1 and LAIR2 expression levels are altered under inflammatory conditions, including infection and autoimmune disease, during activation of T and B cells, and by immune mediators, though the direction of change varies by cell type and condition. For example, activation via the T– or B-cell receptor leads to down-regulation of LAIR1 in T or B cells, respectively, while PMA/ionomycin treatment causes increased secretion of LAIR2 from CD4+ T cells^17–19^. TNFα treatment leads to higher LAIR1 on neutrophils; LPS increases expression of LAIR1 in monocytes and conventional dendritic cells^20, 21^. Cellular location also affects LAIR1 expression levels, as exemplified by neutrophils, in which expression is higher in peripheral blood and lower in bone marrow, spleen, and after extravasation^21, 22^. Though the expression of LAIR genes has been well-studied, relatively little is known regarding the epigenetic and transcriptional regulatory control of these important immune regulators.

Gene regulatory elements, such as enhancers, regulate gene expression in response to cellular and environmental cues, including developmental and immune stimuli. We previously identified enhancers that control the expression of *LAIR2* in normal T cells and T cell lymphomas that were resistant to epigenetic therapies^5^. We next sought to define the gene regulatory elements and immune factors that control LAIR1 and LAIR2 expression in murine and human cells. As we show here using publicly available data, expression of human and murine LAIR1 is highest in myeloid cells, and particularly in macrophages. We next measured *Lair1* mRNA and protein in murine bone marrow (BM) monocytes and BM-derived macrophages and demonstrate that levels are several fold higher in BMDM. *Lair1* expression levels also varied in response to various immune stimuli, with higher levels following type I and type II interferon stimulation and lower levels after LPS stimulation. Lung alveolar macrophages also demonstrated lower levels of LAIR1 surface expression following intra-nasal administration of LPS to mice. In contrast to mice, human monocyte-derived macrophages (MDM) exhibited levels of *LAIR1* and *LAIR2* mRNA that were significantly lower after IFN-γ than after LPS; however, surface protein levels of LAIR1 were consistent with the mouse, with lower levels after LPS relative to IFN-γ treatment. Using data from chromatin immunoprecipitation sequencing (ChIP-seq) of histone modifications and luciferase reporter assays, we identified and tested a set of 37 candidate gene regulatory elements for *Lair1*, *LAIR1* and *LAIR2*. We show that several of these elements exhibit regulatory activity at baseline, and their activity is significantly altered by immune stimuli. Finally, we analyzed transcription factor (TF) ChIP-seq and RNA-seq data to identify a distinct set of TFs for each *LAIR* gene that bind to active regulatory elements and whose expression correlates with LAIR gene expression, suggesting that these factors may exert transcriptional control over *Lair1*, *LAIR1*, and *LAIR2*. Taken together, our findings reveal the epigenetic and immune factors that control the expression of LAIR genes in human and murine cells and provide new insights into how these inhibitory factors are differentially regulated in response to distinct inflammatory stimuli.

## METHODS

### Cell Lines

The human myelogenous leukemia cell line K562 was cultured in Roswell Park Memorial Institute (RPMI) media (Gibco) with 10% fetal bovine serum (FBS; MilliporeSigma), and penicillin-streptomycin 100X(Corning). The human cutaneous T-cell lymphoma cell line HUT78 was cultured in Iscove’s Modified Dulbecco’s Medium (IMDM) (Gibco) with 20% FBS and penicillin-streptomycin 100X. Both were maintained at 37°C and 5% CO_2_ in a humidified incubator.

### Mice

C57BL/6 J WT mice were obtained from the Jackson Laboratory and housed in a pathogen-free environment in the WUSM animal facility. All experimental protocols were approved by Institutional Animal Care and Use Committee of Washington University in St. Louis. Bone marrow was isolated from mice at 7-9 weeks of age.

### Generation of Mouse Bone Marrow-Derived Macrophages

Bone marrow was isolated from mouse femurs and tibias, filtered through a 40-70μm filter (VWR) and cultured in differentiation media consisting of α-Minimal Essential Medium (MEM) (Thermo-Fisher Scientific) containing 10% FBS, 100U/mL penicillin (Corning), 100μg/mL streptomycin (Corning), 50μM β-mercaptoethanol (MiliporeSigma), 2mM L-glutamine, and 10 ng/mL M-CSF from CMG14-12 cells (Veis lab, Washington University in St. Louis) for 7 days with fresh media every three days at 37°C and 5% CO2 in a humidified incubator. For analysis of LAIR1 expression during monocyte to macrophage differentiation from bone marrow, four million undifferentiated bone marrow cells were plated per well of a 6-well plate, then harvested after zero, four and seven days of culture, cells were harvested. For stimulation of BMDMs, whole bone marrow from one mouse was split evenly among the wells of a 6-well plate. For flow cytometry, adherent BMDMs after four or more days of culture were removed by incubation with 0.05% Trypsin-EDTA (Gibco, 25300054) for three minutes at 37°C and 5% CO_2_ followed by forceful pipetting to suspend the cells.

### Generation of Human Monocyte-Derived Macrophages

Human peripheral blood samples were collected from male and female healthy donors aged 20-50 years (2 collections from 3 donors). PBMCs were isolated from blood via density centrifugation using Lymphoprep (StemCell Technologies Cat #07801). The interphase was collected and red blood cells lysed (155 mM NH_4_CL, 10 mM KHCO_3_, 0.1 mM EDTA) for 10 minutes. After a PBS wash, PBMCs were plated at one million cells per mL of RPMI 1640 media with 50ng/mL human M-CSF (M6518, MilliporeSigma), 10% FBS, 100U/mL penicillin (Corning), 100μg/mL streptomycin (Corning), and 50μM β-mercaptoethanol (MiliporeSigma). Cells were differentiated for seven days, with media replaced every three days.

### Cell stimulation

BMDMs were stimulated in complete αMEM with the following proteins as indicated: 20ng/mL mouse IFN-γ (485MI100, R&D Systems), 20ng/mL mouse IFN-β (8234MB010, R&D Systems), 20ng/mL mouse TNF-α (410MT010, R&D Systems), 100ng/mL LPS (L3012, MilliporeSigma), or 50ng/mL IFN-γ and 100ng/mL LPS when used in combination. MDMs were stimulated in complete RPMI with either 10 ng/mL human IFN-γ (285-IF-100, R&D Systems) or 50 ng/mL LPS. K562 cells were stimulated in complete RPMI media with 20 ng/mL human IFN-γ (285-IF-100, R&D Systems), 20 ng/mL human IFN-β (84-99I-F010, R&D Systems), 20 ng/mL human TNF-α (H8916-10UG, MilliporeSigma), 100 ng/mL LPS, or 50 ng/mL human IFN-γ + 100 ng/mL LPS. HUT78 cells were stimulated in complete IMDM media with 20 ng/mL human IFN-γ (285-IF-100, R&D Systems). All treatments were 24 hours.

### Quantitative Reverse Transcriptase-PCR

RNA was directly extracted from macrophages using the RNeasy Plus Mini kit (Qiagen, 74034) and cDNA synthesized using SuperScript™ VILO™ cDNA Synthesis Kit (Invitrogen) following manufacturer’s protocols. SYBR™ Green PCR Master Mix was used to perform transcript quantification. Each reaction was plated in triplicate in a 96-well plate. Relative expression to untreated bone marrow cells or unstimulated BMDMs was calculated via the ΔΔCT method. For specific primers, refer to **Table S1**.

### Mouse LPS treatment and lung cell isolation

Seven week-old mice were anesthetized and treated oropharyngeally with 25 μg LPS (MilliporeSigma, L9143) suspended in 50 μL PBS. After twenty-four hours, bronchoalveolar lavage was performed using 2.5 mL ice-cold PBS supplemented with protease inhibitor (11873580001, MilliporeSigma). Cells were washed with PBS, red blood cells lysed using the same buffer as above for 2 minutes, then washed again prior to staining for flow cytometry. For macrophage gating schema, refer to **Figures S1-S2**.

### Flow cytometric analysis

All incubations were performed at 4 °C in the dark. Isolated cells were incubated with 1:1000 LIVE/DEAD Fixable Violet Dead Stain (L34964, Invitrogen) for 20 minutes. After washing, non-specific Fc binding was blocked with TruStain FcX PLUS (BioLegend) for 10 minutes, then stained with primary antibodies for 30 minutes (see **Table S2**). Analysis was completed with FlowJo software (Version 10.8.1).

### Luciferase reporter assays

To generate luciferase reporter plasmids, candidate regulatory elements (RE) were PCR amplified from human or mouse genomic DNAwith Accuprime Pfx DNA polymerase (Invitrogen) using primers with a NheI or XhoI cut site added to their 5′ end (primer sequences available in **Table S3**). PCR products were digested with NheI and XhoI (New England Biolabs), gel-purified (Qiagen cat. 28,706), and ligated into promoterless pNL1.2[NlucP] or minimal promoter pNL3.2[NlucP/minP] (both from Promega) plasmids using T4 DNA ligase (New England BioLabs, cat. M0202S). Sanger sequencing confirmed successful cloning.

Luciferase reporter assays were performed by electroporation of 1 × 10^6^ K562 or HUT78 cells using the Amaxa Biosystems Nucleofector 2b system with the T-016 or X-001 program setting in Chica nucleofection buffer 1M (5mM KCl; 15mM MgCl2; 120mM Na2HPO4/NaH2PO4 pH7.2; 50mM Mannitol)^23^. Cells were co-transfected with pGL4.54 (transfection control) and luciferase reporter plasmids containing one of the *Lair1*, *LAIR1* or *LAIR2* RE regions. Positive control luciferase reporter plasmids used for cytokine stimulations: pGL4.45 ISRE and pNL3.2.NF-kB-RE, both from Promega. Six hours post-nucleofection, cells were treated with immune stimuli as listed above. Luciferase activity was measured in triplicate per condition in a 96-well plate 24-36 hours post-transfection using the Nano-Glo Dual-Luciferase Reporter System (Promega #N1610) according to the manufacturer’s instructions in a BioTek Cytation5 plate reader. All experiments were performed at least three times. The ratio of the reporter plasmid to the transfection control plasmid luminescence was calculated for each replicate and then normalized for to the average ratio for the empty vector in the matched stimulation condition to determine relative luciferase activity.

### Transcription factor binding site analysis

Transcription factor non-redundant peak sets for hg38 and mm10 were downloaded from the ReMap 2022 website^24^, a database of aggregated publicly available ChIP-seq experiments. Intersection of luciferase regions with transcription factor binding sites was computed with bedtools v2.30.0, then the intersected peak set was filtered in R v4.1.3 using the following criteria: 1) transcription factor has more binding sites in active regulatory elements than in those with no activity (by luciferase reporter) and have more than one binding site across all active regulatory elements; 2) transcription factor is expressed in cell types of interest (splenic red pulp macrophages for *Lair1* (NCBI GEO dataset GSE109125), CD14+ monocytes for *LAIR1*, CD4+ effector T-cells for *LAIR2* (Immgen human PBMCs NCBI GEO dataset GSE227743); 3) transcription factor expression positively correlates with LAIR PBMC expression (using same datasets as in (2)); and 4) transcription factor expression increases following IFN-γ stimulation in cell types of interest (BMDM for *Lair1* (NCBI GEO dataset GSE199128) and *LAIR1*; NK cells for *LAIR2* ^25^). From the resulting set of genes, genes were manually removed if they have DNA-binding activity but no transcription factor activity (RAD21, for example, a component of the cohesion complex) and selected the remaining genes with the highest expression increase following IFN-γ stimulation. R code is available at upon request.

### Publicly available datasets used for transcriptome and epigenome analyses

1. The Human Protein Atlas single cell RNA sequencing dataset; https://www.proteinatlas.org/ ^26, 27^
2. Immgen Mouse ULI Systemwide RNA-seq profiles (NCBI GEO dataset GSE109125) ^28, 29^
3. Immgen Human PBMCs RNAseq (NCBI GEO dataset GSE227743)^30^
4. RNAseq profiling of murine BMDM (NCBI GEO dataset GSE199128)^31^
5. Single Cell Portal; Study: ICA: Blood Mononuclear Cells PBMCs from 2 donors, profiled by SingleCell RNAseq (10X platform) at two ICA sites, (Broad/Boston and MtSinai/NYC).^32^
6. Tabula Muris scRNAseq datasets ^33, 34^
7. ReMap database of Transcription Factor binding^24, 35^

### Statistical analysis

GraphPad Prism 9 (Version 9.4.1) was used for all statistical tests.

## RESULTS

### LAIR1 expression increases as murine bone marrow monocytes differentiate into macrophages in vitro

*Lair1* is expressed in most immune cell subsets, but levels vary widely across subsets, differentiation states, and tissues^10, 16^. Publicly available data demonstrates that *Lair1* mRNA expression is relatively low in lymphocytes and highest in monocytes and macrophages (**Fig. S1A**). Because *Lair1* expression is highest in monocytes and macrophages, we focused on these subsets. Tissue-resident macrophage *Lair1* mRNA expression is high across lung, liver, kidney, brain, spleen, and small intestine (**Fig. S1A** and ^16^), but few comparisons have been conducted of monocytes and macrophages from the same tissue, and to our knowledge, the dynamics of *Lair1* protein and mRNA expression during bone marrow monocyte differentiation into macrophages have not been evaluated.

To test *Lair1* expression over the course of macrophage differentiation, we treated murine bone marrow cells from 7-9 week-old mice over 7 days with M-CSF and measured *Lair1* expression at the level of RNA and protein at days 0, 4, and 7. By qRT-PCR from total cells at each timepoint, we found that compared to day 0, *Lair1* mRNA expression more than doubled after 4 days and nearly doubled again between 4 and 7 days (**Fig. 1A**). Flow cytometry confirmed that the proportion of CD11b+F4/80+ macrophages in the culture increased over time compared to CD11b+F4/80-Ly6C+ monocytes, with macrophages being the majority of cells at days 4 and 7 (**Fig. S1B-D**). LAIR1 surface protein expression increased in CD11b+F4/80+ macrophages by seven-fold between days 0 and 4 and increased an additional 50% between days 4 and 7 (**Fig. 1B-C**). Compared to day 0, LAIR1 surface protein expression increased by two-fold in CD11b+Ly6C+ monocytes after the 7-day exposure to M-CSF, a smaller increase than in macrophages (**Fig. 1C**). Indeed, macrophages produce significantly more LAIR1 than Ly6C+ monocytes at all time points measured (**Fig. 1C**), which is consistent with publicly available expression data comparing *Lair1* RNA expression in mature macrophages and Ly6C+ monocytes (**Fig. 1D**).

**Figure 1.**
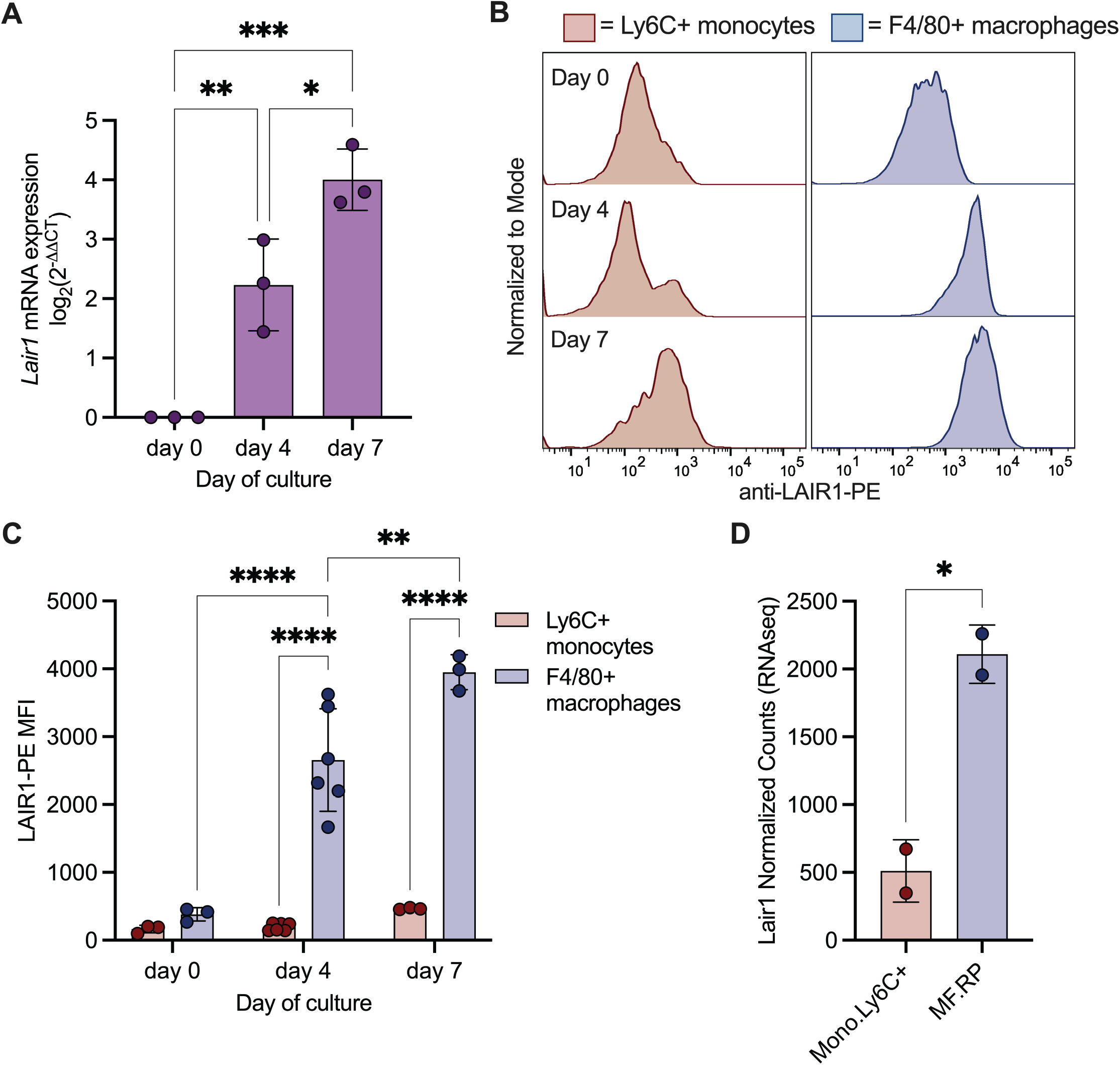
LAIR1 expression increases as murine monocytes differentiate into macrophages from bone marrow *in vitro*. Murine BMDMs were analyzed at three time points during differentiation in culture over 7 days. *Lair1* mRNA expression was measured from total cells by qRT-PCR (**A**) and from monocytes and macrophages by flow cytometry (**B,C**). Representative histograms from each day (**B**) and LAIR1-PE MFI (**C**) are displayed for CD11b+Ly6C+ monocytes (red) and CD11b+F4/80+ macrophages (blue). Statistical significance (two-way ANOVA with Tukey test for multiple comparisons): * p<0.05, ** p<0.01, *** p<0.001, **** p<0.0001. **D**) Publicly available *Lair1* RNA-seq expression data^28, 29^ is plotted as normalized gene counts for Ly6C+ monocytes (Mono.Ly6C+) and splenic red pulp macrophages (MF.RP). Statistical significance (unpaired t-test): * p<0.05

### LAIR1 expression increases in response to interferons and decreases or is unchanged in response to other immune stimuli in mouse BMDM

LAIR1 expression has been shown to increase in response to activating immune stimuli in several immune cell subsets^19–21^, but to our knowledge, murine macrophages have not been evaluated. We used the same model of BMDM (**Fig. 1**) to determine whether LAIR1 expression levels are modulated by immune stimuli in this cellular context. LAIR1 is an inhibitory receptor; thus we hypothesized that early signals in infection (e.g., LPS, TNFα) would cause decreased LAIR1 levels to prevent inhibition of immune response, so-called threshold-disinhibition ^1^. We predicted that signals later in infection (e.g., interferons), would increase LAIR1 expression to down-modulate the immune response and prevent excess damage to the host, termed negative feedback inhibition^1^. We detected slightly decreased levels of *Lair1* RNA in response to 24-hour treatment with TNFα or LPS, with unchanged to lower levels of surface LAIR1, though these changes were not significant. In contrast, we detected a greater than two-fold increase in *Lair1* mRNA (**Fig. 2A**) and an 8-10-fold increase in surface protein (**Fig. 2B-C**) in response to IFN-γ and IFN-β, as we hypothesized. Combining LPS with IFN-γ resulted in decreased levels of mRNA yet slightly increased levels of surface protein. **Figure 2C** shows the distribution of LAIR1 surface expression across BMDM treated with different stimuli from a representative experiment. Compared to baseline, which has a relatively broad distribution of LAIR1 expression across the macrophage population, TNFα, LPS, and LPS with IFN-γ maintained the distribution size with minor shifts in geometric mean, indicating that there is a range of surface LAIR1 across the population. In contrast, IFN-γ and IFN-β treatment cause a narrowing of the distribution with a rightward shift, indicating that all macrophages have increased their surface LAIR1 expression. Overall, these findings are consistent with our hypothesis, though increased LAIR1 expression in response to interferons was significantly more robust than were decreases in response to TNFα and LPS. To further evaluate LAIR1 expression in response to immune stimuli, we next employed an *in vivo* model.

**Figure 2.**
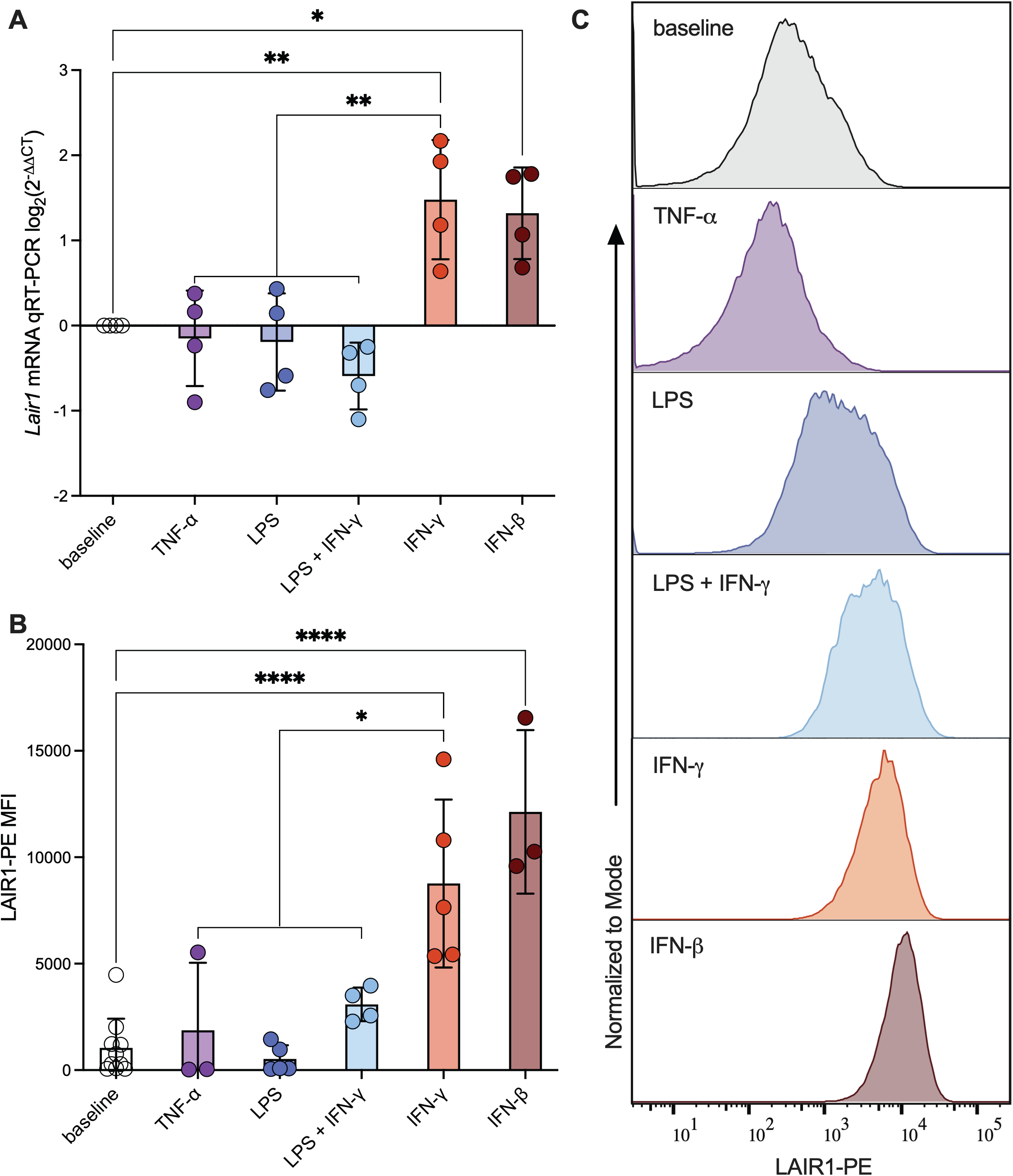
LAIR1 expression increases in response to interferons and decreases or is unchanged in response to other immune stimuli in mouse BMDM. Murine BMDMs were differentiated in culture for 7 days, then treated with IFN-γ, IFN-β, TNF-ɑ, LPS, LPS + IFN-γ, or were left untreated for 24 hours (n=3-8). *Lair1* mRNA expression was measured from total cells by qRT-PCR (**A**) and from CD11b+F4/80+ macrophages by flow cytometry (**B,C**). LAIR1-PE MFI (**B**) and representative histograms (**C**) are shown for CD11b+F4/80+ macrophages from each condition. Statistical significance (one-way ANOVA with Tukey test for multiple comparisons):* p<0.05, ** p<0.01, *** p<0.001, **** p<0.0001

### In vivo LPS treatment downregulates LAIR1 expression in mouse alveolar macrophages

After determining that *Lair1* expression level changes in response to immune stimuli in BMDM, we next sought to test whether these changes also occur in macrophages *in vivo.* We decided to focus on lung tissue-resident macrophages due to high *Lair1* expression in alveolar macrophages (AVM) overall and compared to other immune cell subsets in lung, and increased *Lair1* RNA expression on AVMs in a lung tumor model (**Fig. 3A, Suppl. Fig. 1A** and ^16^). Oropharyngeal LPS administration in mice is a well-characterized model of acute lung inflammation and allowed us to examine a stimulus *in vivo* that produced decreased *Lair1* expression in BMDM *in vitro* (**Fig. 2, 3B** and ^36^). We therefore treated wild-type C57/Bl6 mice with PBS (vehicle) or LPS via oropharyngeal inoculation and collected bronchoalveolar lavage fluid (BALF) 24 hours later. BALF cells were stained for CD45, CD11c, SiglecF, and LAIR1 to identify AVMs (CD45+CD11c+SiglecF+) and measure LAIR1 expression using flow cytometry (**Fig. S2**). We isolated similar numbers of AVMs from PBS– and LPS-treated mice, which is expected in this model^36^ (**Fig. 3C**). Consistent with our hypothesis and *in vitro* studies, LAIR1 surface expression decreases by more than 30% on AVMs after LPS treatment (**Fig. 3D-E**).

**Figure 3.**
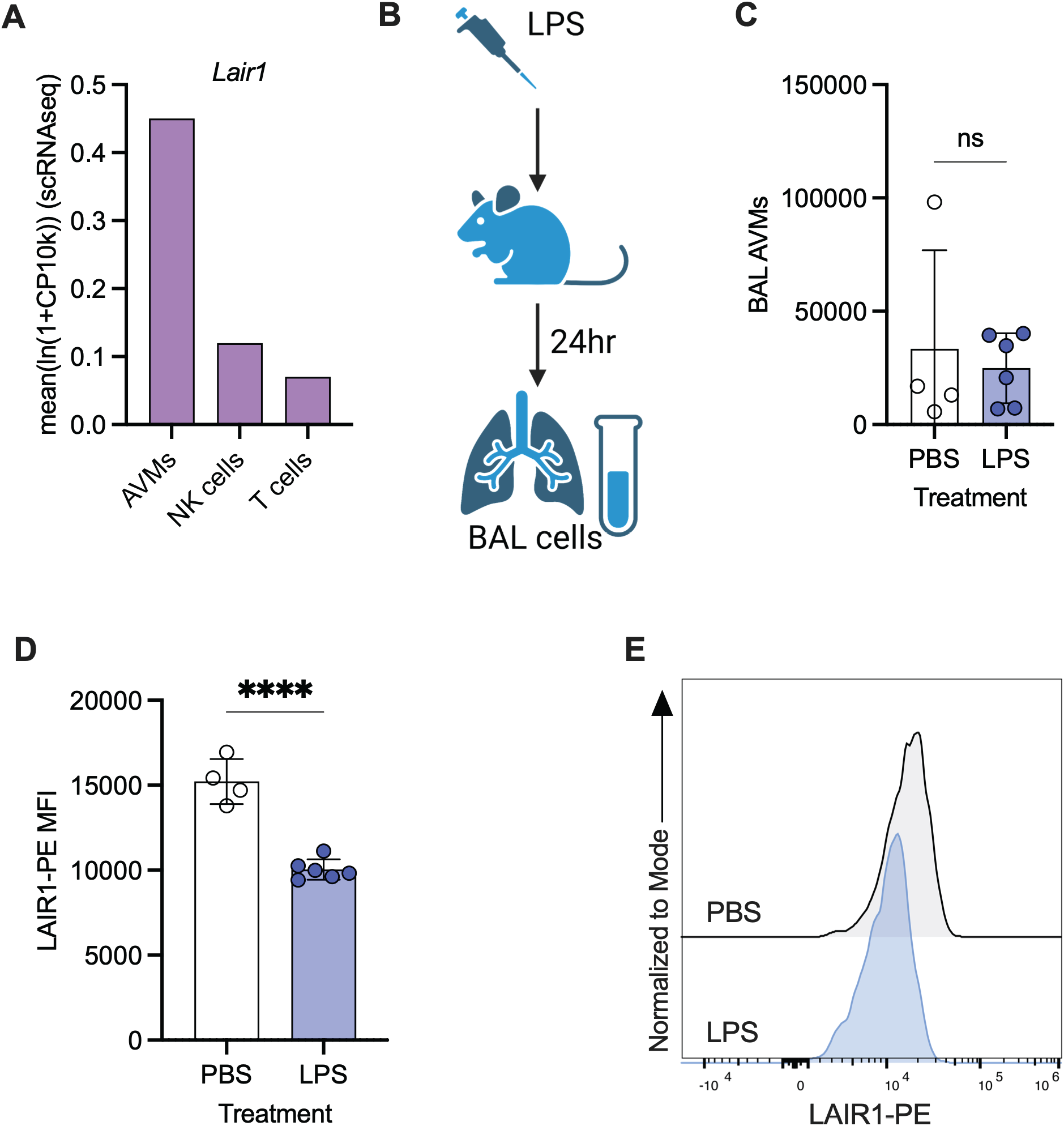
LAIR1 expression in mouse alveolar macrophages decreases after *in vivo* LPS treatment. **A**) Publicly available *Lair1* expression data from the Tabula muris droplet lung dataset^33^ plotted for alveolar macrophages (AVMs), natural killer cells (NK cells), and T-cells. Bronchoalveolar lavage cells were collected from wild type mice (n=4-6) 24 hours after oropharyngeal inoculation with PBS or LPS (**B**), then analyzed by flow cytometry. Total number of AVMs (CD45+CD11c+SiglecF+) isolated (**C**), LAIR1-PE MFI (**D**), and representative histograms (**E**). Statistical significance (unpaired t-test): **** p<0.0001

### LAIR1 and LAIR2 expression patterns are distinct from Lair1

In contrast to mouse, the human genome contains both *LAIR1* and *LAIR2* genes, which arose due to a genomic duplication inversion event^14^. These genes have non-overlapping expression patterns across immune cell subsets, with generally higher expression of *LAIR1* than *LAIR2* in myeloid subsets. Indeed, single cell RNAseq data demonstrates that *LAIR1* expression is over 100-fold higher in macrophages than *LAIR2*, while the latter is relatively higher in other cell types, including NK and T cells (**Fig. 4A, S3A-B** from the Human Protein Atlas^26, 27, 32^ and Single Cell Portal, see Methods). Previous reports differ in the impact of immune stimuli on *LAIR1* expression in human monocytes, with both increased and decreased surface expression detected after treatment with microbial or inflammatory factors^20, 37^. Treatment of monocytes for seven days in media containing immune mediators resulted in a range of monocyte/macrophage phenotypes and variable levels of LAIR1^20^. We therefore sought to determine the impact on *LAIR1* expression of treating M-CSF-differentiated monocyte-derived macrophages (MDMs) with short-term immune stimuli.

**Figure 4.**
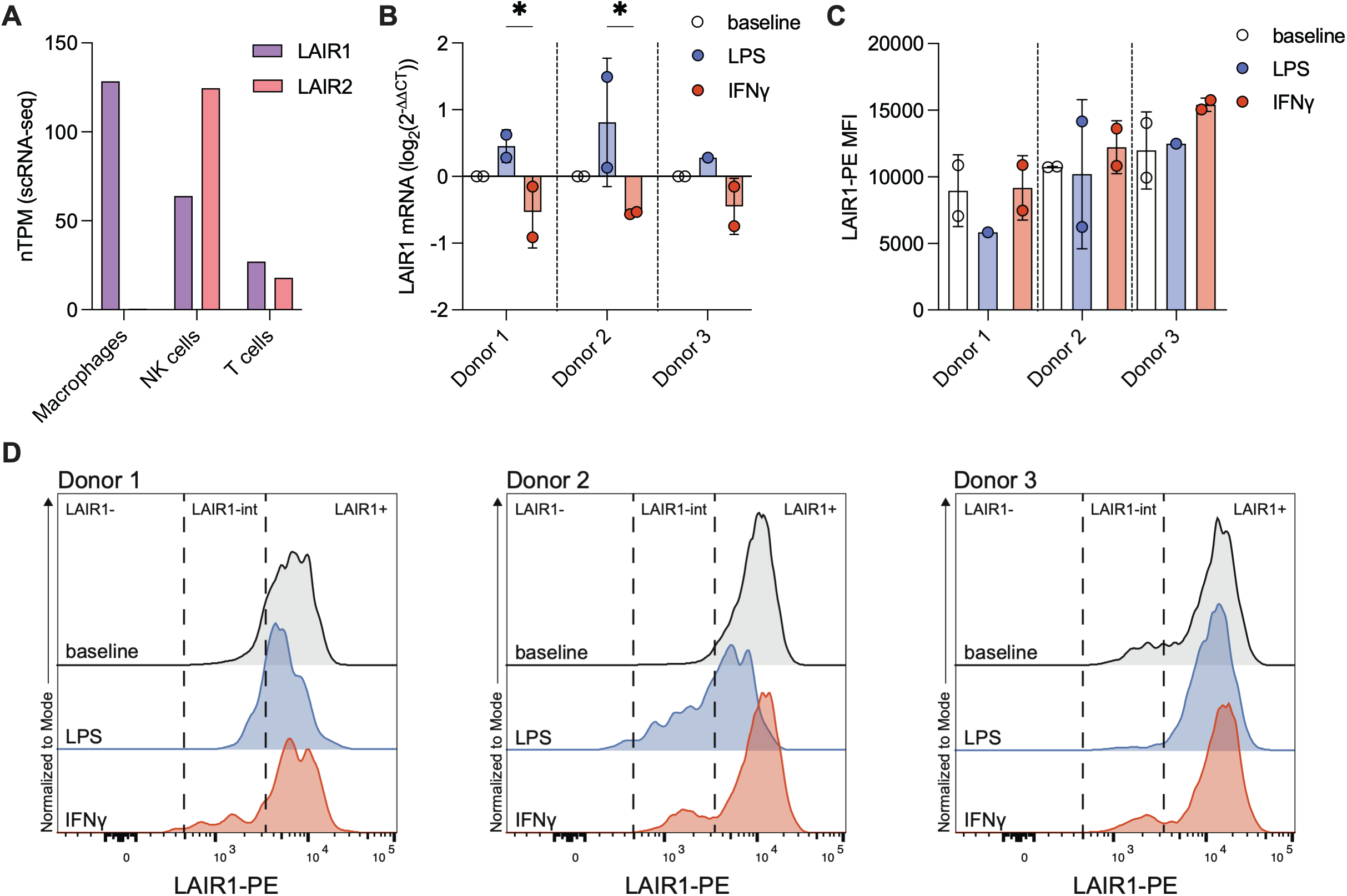
LAIR1 and LAIR2 expression patterns are distinct from Lair1. PBMCs were isolated from three healthy donors and differentiated over 7 days to form MDMs. **A**) *LAIR1* and *LAIR2* expression (scRNAseq, Human Protein Atlas^26, 27^. **B**) cDNA was isolated from MDM day 7 cultures after 24 hours IFNγ or LPS stimulation and qRT-PCR for *LAIR1* was performed. Relative expression was measured as the inverse of dCt. **C**) LAIR1 surface protein expression was measured by flow cytometry in MDMs (CD11b+CD68+) after 24 hours IFNγ or LPS stimulation. **D**) Histograms from a representative experiment demonstrating LAIR1 protein expression in individual donors following 24 hours IFNγ or LPS stimulation. Statistical significance (one-way ANOVA with Fisher’s LSD test): *p<0.05

To address this question, we isolated peripheral blood cells from three healthy donors and differentiated total PBMCs into monocyte-derived macrophages (MDMs) by treatment with human M-CSF for 7 days. Flow cytometry demonstrated that nearly 80% of cells were CD11B+CD68+ MDMs (**Fig. S3C**). Due to limiting cell numbers, we were able to test fewer conditions, thus we chose stimuli that caused markedly different *Lair1* expression in mouse macrophages, LPS and IFN-γ (**Fig. 2**). As expected, MDMs from 3 human donors showed greater variability in their baseline and stimulated levels of *LAIR1* expression compared to BMDMs from inbred mice, but consistent patterns emerged (**Fig. 4B-D, Table S4**). Total day 7 cells treated for 24 hours of with LPS had *LAIR1* mRNA levels that were consistently and significantly higher than after 24 hours of IFN-γ treatment. In contrast to mRNA and consistent with the mouse, surface LAIR1 levels were lower in LPS-treated than IFN-γ-treated CD11B+CD68+ MDMs (**Fig. 4C-D**). These data may reflect timing differences in transcriptional versus protein level changes or may be due to the presence of ∼20% non-macrophage cells in the mRNA analysis (**Fig. S3C**). *LAIR2* exhibited much lower mRNA expression levels than *LAIR1* in MDMs (**Table S4**), consistent with publicly available data (**Fig. 4A, S3A**). *LAIR2* mRNA levels varied by donor, but were overall less affected by LPS than *LAIR1* (**Fig. S3D**). Conversely, after 24 hours IFNγ stimulation, *LAIR2* mRNA levels were decreased in all donors (**Fig. S3D**). Overall, changes in LAIR expression in response to immune stimuli were markedly different in human and mouse macrophages, though LAIR1 surface protein was consistently lower in LPS-treated compared to IFN-γ-treated macrophages.

### Identification of candidate enhancers and promoters

The differences in response to immune stimuli in mouse and human macrophages led us to hypothesize that they might be due, at least in part, to differences in the transcriptional regulatory elements (enhancers, promoters) that control LAIR transcription. We used publicly available histone modification chromatin immunoprecipitation sequencing (ChIP-seq) and chromatin accessibility (ATAC-seq) datasets (ENCODE^38, 39^), ReMap transcription factor binding density (TF-ChIP-seq) datasets^24^, and our previously published histone modification ChIP-seq from a Cutaneous T-Cell Lymphoma (CTCL) sample with altered enhancer activity and increased expression of LAIR genes^5^ to identify candidate enhancers and promoters for mouse *Lair1* and human *LAIR1* and *LAIR2*. Certain combinations of histone modification peaks detected by ChIP-seq indicate that a genomic region may be a transcriptional regulatory element. Specifically, active promoter and enhancer elements have regions of open chromatin (ATAC– or DNAse-seq peaks), and may also have H3K9ac, H3K27ac, and TF-ChIP-seq peaks. In addition, promoters typically harbor H3K4me3, while active enhancer elements have H3K4me1 modifications^40–42^.

Applying these criteria to mouse ChIP– and ATAC-seq data from the mouse *Lair1* gene and flanking regions, we identified one candidate promoter region and six upstream candidate enhancer elements that largely overlapped with ENCODE^38, 39^-predicted elements (cCREs) and harbored H3K4me1, H3K27ac, H3K9ac peaks, increased chromatin accessibility, and increased TF binding density (**Fig. 5A-B**). For the human *LAIR1* (**Fig. 5C**) and *LAIR2* (**Fig. 5E**) genes and flanking regions, we applied the same criteria to ENCODE ChIP– and DNAase-seq data from many cell types, including the myeloid cell line K562, as well as H3ac and H4K27ac ChIP-seq from T-cells from our previous study^5^, and identified 9 candidate enhancers and 2 candidate promoters for *LAIR1* (**Fig. 5D**) and 13 candidate enhancers and 2 candidate promoters for *LAIR2* (**Fig. 5F**). These chromatin patterns are quite similar in primary human CD14+ monocytes and CD4+ αβ T cells, with higher levels of activating histone marks in monocytes, as expected (**Fig. S4A-B**). Two of the *LAIR2* candidate regulatory elements (h18 and h19) overlap with regulatory elements that we previously identified in HDACi-resistant CTCL. *LAIR1* and *LAIR2* genes each have alternative transcript isoforms in the ENSEMBL database that are similar to canonical/default transcripts but have alternative first exons and transcriptional start sites (TSS) that are ∼5kb upstream of the canonical TSS; these coincide with candidate regulatory elements h6 and h19 (**Fig. 5D, F**). For *LAIR1*, the genomic location of the alternative first exon coincides with an ENCODE candidate promoter element, and for both *LAIR1* and *LAIR2*, these candidate alternative promoters have H3K4Me3 peaks. RNAseq data from CD14+ monocytes and CD4+ T cells shows expression peaks in the upstream alternative exons and TSS of *LAIR1* and *LAIR2*, with higher expression in these regions in monocytes (**Fig. S4A-B**). In summary, we identified 37 candidate promoter and enhancer regulatory elements for mouse and human LAIR genes that may control their transcription under homeostatic and stimulated conditions.

**Figure 5.**
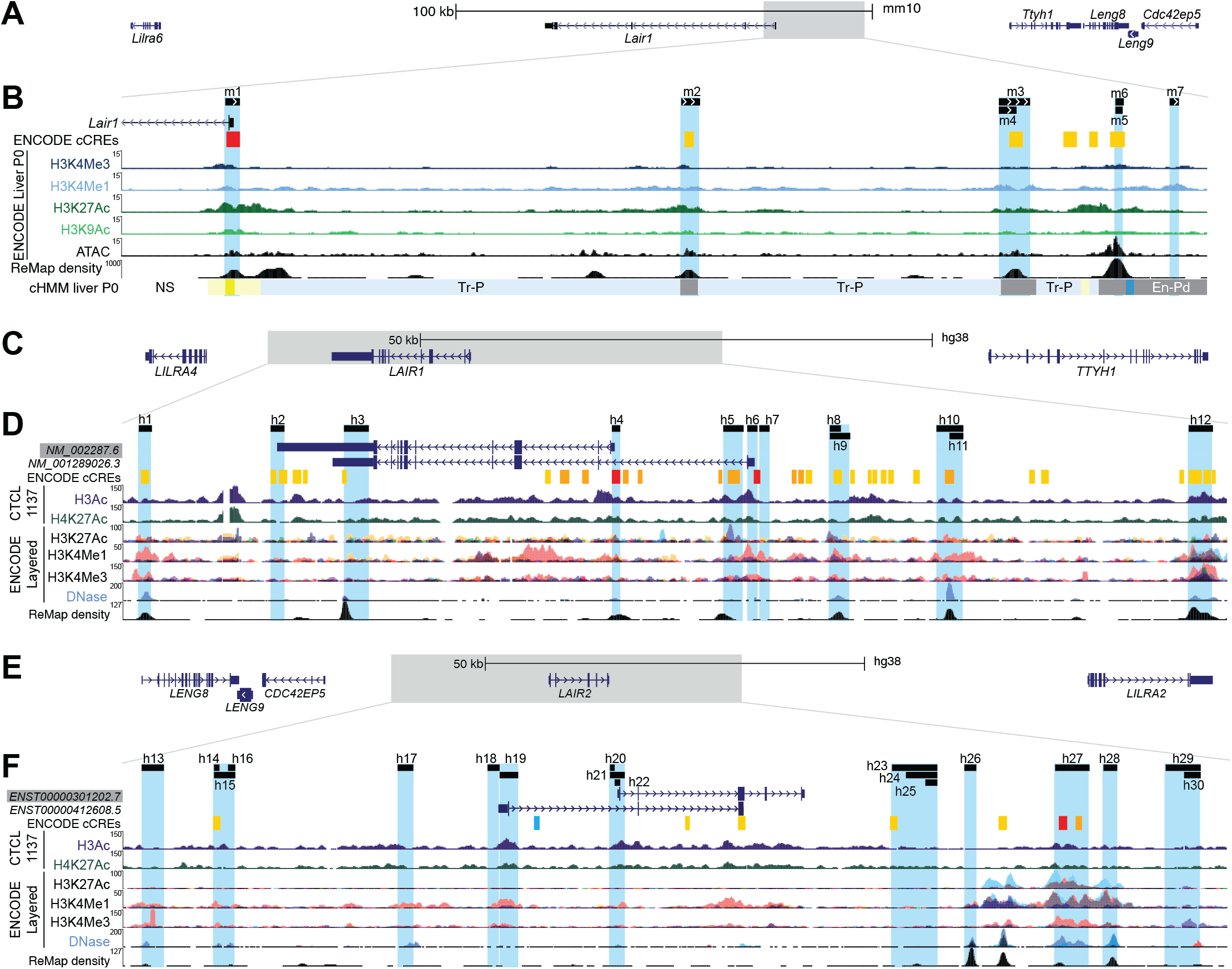
Candidate regulatory elements identified for *Lair1*, *LAIR1* and *LAIR2*. UCSC genome browser tracks^87^ showing mouse *Lair1* (mm10, Jan. 2012) and human *LAIR1* and *LAIR2* (hg38, Dec. 2013). For each gene, nearest neighbor genes and regions tested for regulatory activity (whole area highlighted in gray) are shown (**A, C, E**). Zoomed in views for each gene with candidate regulatory elements tested in reporter assays labeled and highlighted in blue (**B, D, F**). The *Lair1* track is shown with ENCODE^39, 87^ tracks for candidate cis-regulatory elements (cCREs, red = promoter, yellow = enhancer); postnatal day 0 (P0) liver ENCODE ATAC-seq signal and histone modification ChIP-seq signal (H3K4Me3, H3K4Me1, H3K27Ac, H3K9Ac); ReMap ChIP– seq^24^ total transcription factor binding density for BMDMs; and ENCODE ChromHMM chromatin state from P0 liver (**B**). *LAIR1* & *LAIR2* tracks are shown with ENCODE cCREs (red = promoter, yellow = enhancer); histone modifications (H3Ac, H4K27Ac) for CTCL patient 1137^5^; ChIP-Seq histone modifications from ENCODE Chip-seq layered 7 cell lines (H3K27Ac, H3K4Me1, H3K4Me3); ENCODE DNaseI hypersensitivity signal from 9 cell types; and ReMap ChIP-seq total transcription factor binding density (**D, F**).

### Regulatory element activity is altered by immune stimuli in distinct patterns in human and mouse LAIR genes

Having identified these 37 candidate regulatory elements for mouse and human LAIR genes, we next sought to test whether they harbor functional gene regulatory activity. We subjected the candidate elements to luciferase reporter assays by cloning them into the promoter site of a promoterless reporter or the distal (enhancer) site of a minimal promoter reporter. We next transfected them into the myeloid cell line (K562), treated with immune stimuli as in **Figs. 2** and **4**, and measured transcriptional regulatory activity using protocols adapted from our previous work^5^. Because all mouse *Lair1* transcripts have the same promoter (**Fig. 5A**), we tested all mouse candidate elements using the enhancer reporter. As expected, the *Lair1* promoter element (m1) did not show enhancer activity at baseline or with immune stimuli; the lack of activity in the m7 candidate enhancer element also was not surprising given that it harbored only the poised enhancer mark H3K4me1 and no activating chromatin marks or TF binding peaks (**Fig. 5B, 6A, S5A**). Two other candidate elements (m2, m4) showed minimal to no activity, despite having high levels of active chromatin marks and TF binding (**Fig. 5B, 6A**). In contrast to these, several candidate enhancers that lie 20kb upstream of the *Lair1* promoter exhibited significantly higher levels of transcriptional regulatory activity compared to control (empty vector) (**Fig. 6A**). One candidate enhancer (m6) exhibited five-fold higher baseline activity (i.e., without immune stimulation) compared to control; after treatment with immune stimuli, m6 and two others (m3, m5) exhibited significantly higher activity compared to baseline. All three candidate enhancers exhibited a two-fold or greater increase in activity compared to baseline after IFN-γ, IFN-β, TNF-α, and LPS+IFN-γ. Both m3 and m6 also exhibited at least two-fold increased luciferase activity after LPS alone, whereas m5’s activity was not significantly higher after LPS alone. In summary, treatment with IFN-γ (m3, m6) and IFN-β (m5) caused the greatest increase in regulatory activity for these enhancer elements, which is consistent with our data showing that the largest increases in *Lair1* expression occurred after treatment with IFN-γ or IFN-β (**Fig. 2**).

**Figure 6.**
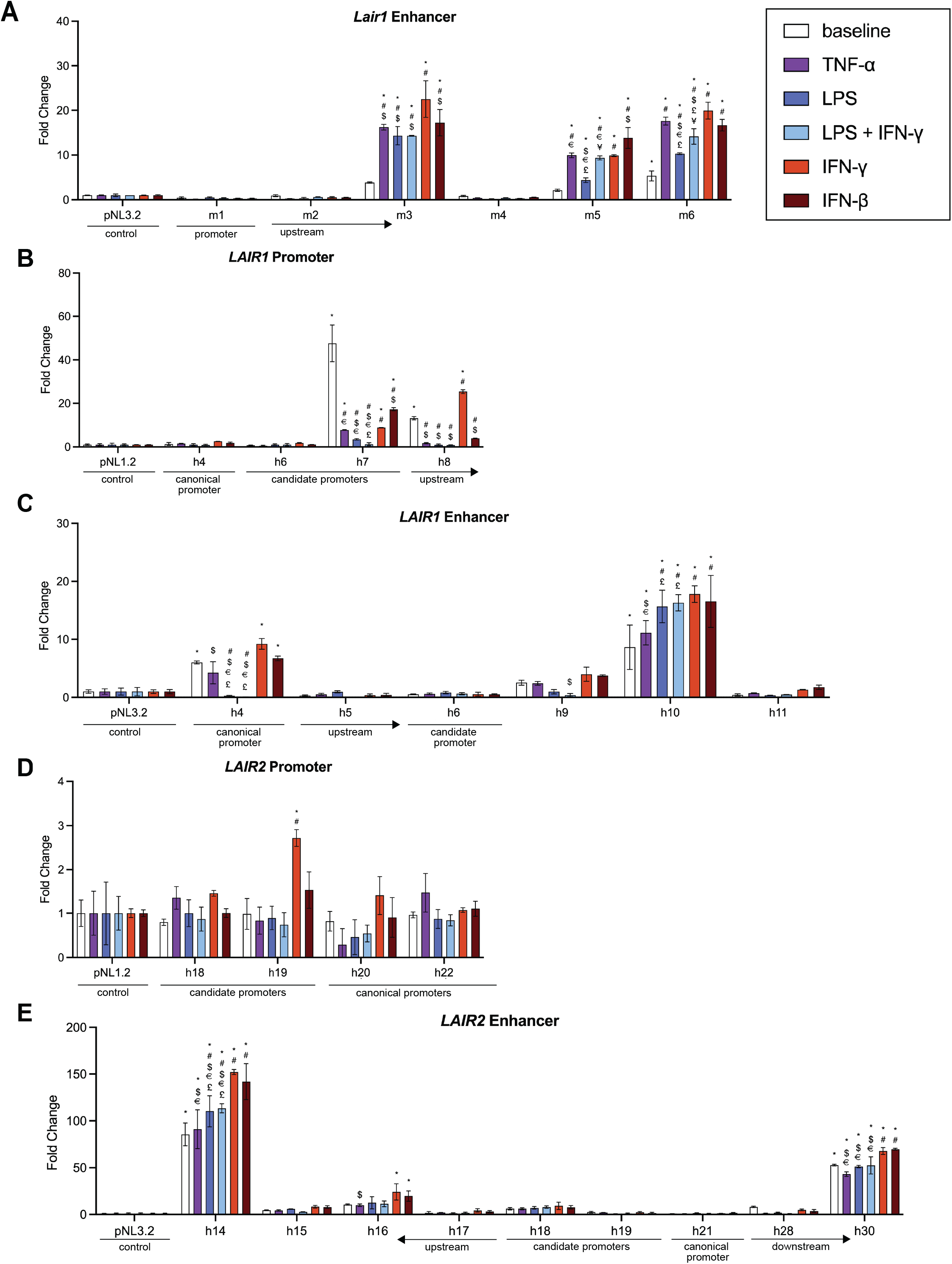
LAIR candidate distal enhancers exhibit high baseline activity that is modulated following immune stimuli. Bar graphs show fold change and standard deviation for candidate regulatory elements for *Lair1* enhancer reporter (**A**), *LAIR1* promoter and enhancer reporters (**B-C**), and *LAIR2* promoter and enhancer reporters (**D-E**) relative to empty vector control in K562 cells after 24 hours of indicated stimulations. Statistical significance (two-way ANOVA with Tukey’s test for multiple comparisons) shown for a subset to facilitate ease of comparison: * p<0.05 for condition compared to its empty vector control; # p<0.05 for immune-stimulated condition compared to baseline for same candidate region; $ p<0.05 for condition compared to IFN-gamma-stimulated RE; € p<0.05 for condition compared to IFN-β-stimulated RE; £ p<0.05 for condition compared to TNFα-stimulated RE; ¥ p< 0.05 for condition compared to the same LPS-stimulated RE. For a full list of statistical comparisons, refer to **Table S5**.

The human *LAIR1* locus is similar to that of mouse *Lair1*, in that both genes are on the negative strand, there are large non-coding regions 15-20kb 5’ of the promoters that contain several ENCODE cCREs, and the nearest upstream coding genes are the homologous *TTYH1* and *Ttyh1*, respectively (**Fig. 5A,C).** One difference is that *LAIR1* has an alternative transcript with a transcriptional start site (TSS) ∼5kb upstream of the canonical TSS that overlaps an ENCODE cCRE promoter (**Fig. 5D**). Strikingly, this upstream promoter element (h7) exhibited the highest baseline transcriptional regulatory activity of all elements tested in the promoter reporter, nearly 50-fold higher than control (empty vector) (**Fig. 6B**). The activity of the h7 element was significantly reduced by treatment with all immune stimuli we tested, potentially suggesting that transcription of this alternative transcript would be reduced under inflammatory conditions. An adjacent region (h6) containing the alternative TSS/1^st^ exon had no activity. Surprisingly, the canonical promoter region (h4) had no promoter activity under any condition (**Fig. 6B**). We also detected promoter activity 15-fold higher than control in candidate element h8 that was significantly increased by IFN-γ stimulation but decreased by all other immune stimuli tested. Further evaluation of h8 revealed a lncRNA gene (*ENSG00000288742*), which may be an enhancer RNA (eRNA) based on overlapping enhancer marks and ENCODE designation as a cCRE enhancer (**Fig. 5D**). While the canonical *LAIR1* promoter (h4) did not show activity in the promoter reporter, in the enhancer reporter it showed significantly higher activity compared to empty vector (**Fig. 6C**). Similar to candidate promoter element h7, h4 enhancer activity was significantly reduced by treatment with LPS or LPS + IFNγ, but IFNγ or IFNβ alone did not alter activity from baseline. The h9 element, which encompasses h8, showed a similar pattern of regulatory activity to h8 (**Fig. 6B**), though at lower levels and only the decrease after LPS + IFN-γ treatment was significant (**Fig. 6C**). Distinct from the other *LAIR1* regulatory elements, a candidate enhancer element ∼14kb upstream of the canonical TSS, h10, exhibited significantly increased activity relative to baseline in response to immune stimuli (**Fig. 6C, S5B**), which was similar, though more modest, to the *Lair1* m3 enhancer (**Fig. 6A**). Overall, *LAIR1* regulatory elements had distinct baseline activity levels and responses to immune stimuli that, except for h10-m3, were substantially different from *Lair1*.

For *LAIR2*, we did not detect baseline promoter activity in any element; only one element, h19, which overlaps the alternative upstream TSS, showed increased activity after treatment with IFN-γ (**Fig. 5F, 6D**). The less than three-fold change was considerably smaller than the *LAIR1* candidate promoters, which is consistent with lower expression levels of *LAIR2* in myeloid cells. We next tested 16 candidate regulatory elements for enhancer activity, including five upstream (h13 – h17), three candidate/canonical promoter (h18, h19, h21), and eight downstream (h23-h30) elements, of which two (h14, h30) exhibited activity significantly above baseline (**Fig. 6E**, **S5C**). H14, h16, and h30 also showed significantly altered enhancer activity in response to several immune stimuli, including IFN-γ (**Fig. 6E**, **S5C**). Because *LAIR2* expression levels are higher in T cells than myeloid cells, we tested a subset of regulatory elements in HUT78, a CTCL cell line, including candidate enhancer elements (h14, h16) and candidate promoter elements (h18, h19). H14 and h16 showed a significant 10-fold increase in regulatory activity after treatment with IFN-γ. Interestingly, interferon-stimulated activity levels were similar in h14 and h16 in T-cells, whereas h14 activity was substantially higher than h16 in myeloid cells (**Figs. 6E, S5C-D**). These results are consistent with higher levels of *LAIR2* expression in T cells under inflammatory conditions, such as in autoimmune diseases and CTCL^5, 18^.

In summary, we confirmed regulatory activity in a subset of candidate elements identified by chromatin profiling. We determined that candidate upstream promoter elements (h7, h19) exhibited higher regulatory activity than the canonical promoters. In addition, seven candidate elements exhibited baseline regulatory activity significantly increased compared to control (m6, h4, h7, h8, h10, h14, h30). All seven of these, plus five others in which baseline activity was not different from control (m3, m5, h9, h16, h19), exhibited significant changes in activity, both higher and lower, after immune stimulation. The largest increases were induced by IFN-γ and the largest decreases were induced by LPS. Together, these results suggest that transcriptional regulators activated by the immune stimuli may control expression of *Lair1*, *LAIR1*, and *LAIR2* genes.

### Transcription factors in inflammatory pathways bind active regulatory elements and correlate with LAIR expression

Having shown that expression of LAIR genes and the activity of their transcriptional regulatory elements have distinct patterns in myeloid cells and T cells and are responsive to immune stimuli, particularly after IFN-γ treatment, we next sought to identify the transcription factors that control these changes. We used publicly available TF ChIP-seq data^24^ to identify TF binding sites that overlapped with highly active regulatory elements, and RNA-seq data from monocytes, macrophages, T and NK cells treated with IFN-γ to determine expression patterns of these TF and LAIR genes. We required that: 1) there were more peaks for a transcription factor in highly active regulatory elements than in less active regulatory elements; 2) TF expression correlated with LAIR expression; 3) the TF was expressed in monocytes (*LAIR1*), macrophages (*Lair1*), T or NK cells (*LAIR2*); and 4) that the TF expression increased after IFN-γ treatment. Panels **A-C** of **Figure 7** show a separate track for each TF meeting these criteria; colored blocks mark TF binding site peaks, with wider blocks indicating a higher number of peaks. There are shared and distinct TFs for each LAIR gene. *Lair1* regulatory elements are bound by several factors, including IRF2, MAF, NR3C1, STAT3 AND STAT5B, of which NR3C1 and STAT5B are co-regulators. *Lair1* expression and enhancer element activity were increased in the three active elements (m3, m5, and m6) after most immune stimuli, including IFN-γ, while LPS treatment caused relatively lower expression and reporter activity in m5 and m6 (**Figs. 2, 3, 6A**). Comparison of the TF binding profiles of these elements reveals that m5 and m6, but not m3, harbor TFs known to suppress transcription downstream of LPS stimulation, STAT3 and IRF2^43, 44^ (**Fig. 7A**). *LAIR1* regulatory elements are bound by BACH1, MAFK, PML, RELA and STAT5A, of which BACH1 and MAFK are co-regulators. The variability in *LAIR1* regulatory element activity, in which some elements showed decreases in response to LPS (h4, h7, h8) and IFN-γ (h7), and others showed increases (h8 to IFN-γ, h10 to all immune stimuli) (**Fig. 6B-C**), could be due to distinct TF binding profiles at each site. Notably, h10 lacks binding of BACH1, a known transcriptional suppressor after LPS exposure^45, 46^ (**Fig. 7B**). *LAIR2* regulatory elements are bound by ATF2, JUNB, STAT5A, TBX21 (T-bet), and TRIM22, of which ATF2 and JUNB are co-regulators. *LAIR2* regulatory elements h14 and h16, which have the highest activity in T cells, harbor peaks for the same TFs as other elements with less activity, but have a substantially greater number of peaks bound by JUNB, TRIM22, and TBX21 (T-bet), which are important in T cell activation and inflammatory responses^47–49^. *LAIR1* and *LAIR2* loci each have some regulatory elements with relatively high levels of activity based on chromatin modifications and luciferase reporter assays, but no TF peaks that met our criteria (e.g., h7, h19, h30). This could be due to chromatin modifiers not captured in the TF analysis (e.g., P300, CTCF, HDAC) and/or to 3D interactions with regulatory elements that do have TF binding. Indeed, capture Hi-C genome conformation data^50^ shows genome interactions between most of the regulatory elements, including some of those without TF peaks (**Fig. S6A-B)**. It is interesting to note that the alternative upstream TSS/promoter elements of both *LAIR1* and *LAIR2* have the greatest number of 3D interactions in each gene locus. Finally, the role of these TFs in regulating cell-specific LAIR expression is further supported by higher TF expression levels (darker colors) and larger numbers of TF binding site peaks (larger circles) in highly active regulatory elements in myeloid (*Lair1*, *LAIR1*) or lymphoid (*LAIR2*) cell subsets (**Fig. 7D-F**).

**Figure 7.**
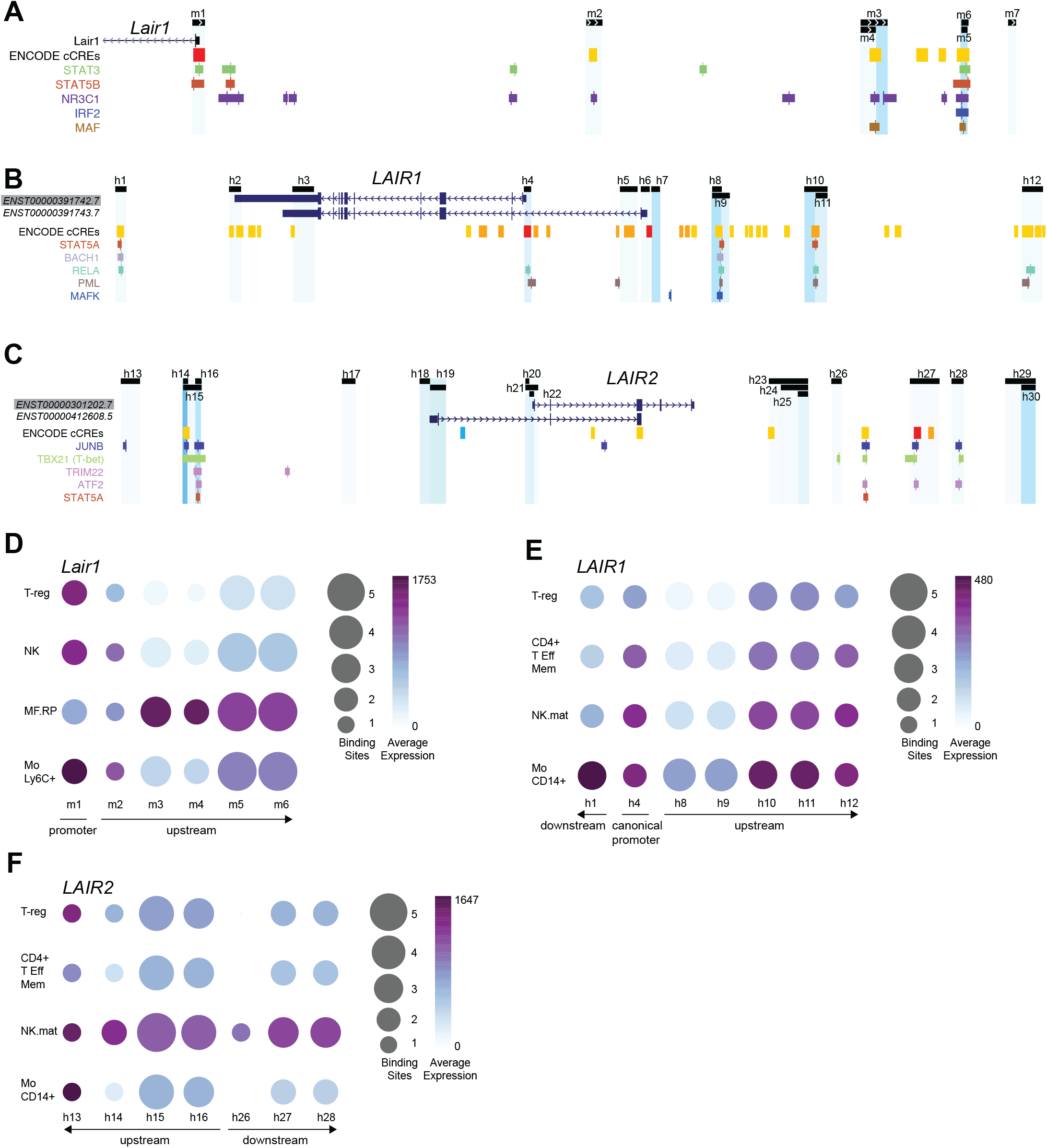
Transcription factors in inflammatory pathways bind active regulatory elements and correlate with LAIR expression. Candidate regulatory elements, ENCODE cCREs^39, 87^, and ReMap ChIP-seq^24^ binding sites for selected transcription factors are shown for *Lair1* (**A**), *LAIR1* (**B**), and *LAIR2* (**C**) over the same genomic coordinates as in Fig. 5B**,D**,**F**. Candidate regulatory elements are highlighted in colors in proportion to maximal luciferase reporter activity (darker indicates higher activity). **D-F**) TF expression level (shown by color, fraction maximal for each cell type) and TF binding sites (shown by circle size) from **A-C** are plotted by selected cell type in mouse (**D**) and human (**E-F**).

## DISCUSSION

LAIR proteins comprise their own small branch of the Ig-like inhibitory immune receptor family^14^. Though LAIR1 and LAIR2 have been separately studied for their transcriptional response to immune stimuli, they have been infrequently examined together. Moreover, the epigenetic and transcriptional control mechanisms that differentially regulate the expression of *Lair1*, *LAIR1*, and *LAIR2* have not previously been characterized.

To address this knowledge gap, we examined gene regulatory control under homeostatic and immune stimulated conditions in mouse and human monocytes, macrophages, and T cells. Previous reports showed that inflammatory stimuli, including autoimmune disease, infection, and immune mediators, alter the levels of LAIR1 and LAIR2 protein in neutrophils, monocytes, and T cells^17, 18, 20, 21, 51, 52^. Due to macrophages’ critical role in both pro-inflammatory immune defense and anti-inflammatory infection resolution, we hypothesized that transcriptional control of LAIR genes may be distinct in these cells. In addition, we sought to compare the gene regulatory landscapes of *Lair1*, *LAIR1* and *LAIR2*. Because LAIR1 is an inhibitory receptor, we predicted that early signals in infection (e.g., LPS, TNFα) would cause decreased LAIR1 levels to prevent inhibition of immune response, a functional distinction that has been termed threshold-disinhibition^1^. On the other hand, we predicted that signals later in infection (e.g., interferons), would increase LAIR1 expression to down-modulate the immune response and prevent excess damage to the host, which has been called negative feedback inhibition^1^. As we show here in macrophages, and as previously reported in other cells, these patterns of response to immune activation are dependent on cellular context and immune stimulus^9, 17, 18, 20, 21, 51, 53^.

Using murine BMDM and human MDM cultures, we measured LAIR transcript and protein during differentiation and after treatment with type I and type II interferons, LPS, and TNFα. Our studies confirmed that expression of LAIR1 increases with macrophage differentiation in culture in both human MDM and mouse BMDM. Work presented here also revealed differences in the levels of *Lair1* and *LAIR1* mRNA and protein in mouse and human macrophages in response to immune stimuli. Of note, *Lair1* expression in macrophages was consistently higher following type I and type II interferon stimulation and lower after LPS stimulation, both *in vitro* and *in vivo*. Combining LPS with IFN-γ resulted in decreased levels of RNA yet slightly increased levels of surface protein, suggesting either a lag in the impact on protein levels or disparate effects of this combination on pathways that control transcript and protein levels. These results are consistent with both threshold inhibition (in response to LPS) and negative feedback inhibition (in response to interferons)^1^.

In contrast to mice, human MDMs exhibited levels of *LAIR1* and *LAIR2* mRNA that were lower after IFN-γ than after LPS; however, surface protein levels of LAIR1 were consistent with the mouse, with lower levels after LPS relative to IFN-γ treatment. These data may reflect timing differences in transcriptional versus protein level changes. Nevertheless, IFN-γ treatment did not increase *LAIR1* levels to the degree that it did for *Lair1*. Together, these results suggest that the transcriptional regulatory mechanisms controlling expression of *Lair1*, *LAIR1*, and *LAIR2* are distinct, which may be due to differences in the epigenetic and transcriptional regulatory landscape of each gene.

To address this question, we assessed the chromatin landscape using histone modification and transcription factor ChIP-seq and ATAC-seq to define 37 candidate gene regulatory elements and transcription factors, then tested these elements in luciferase reporter assays. In total, 12 elements had regulatory activity at baseline that was significantly different from control and/or was significantly changed by immune stimuli. Overall, the murine *Lair1* locus contained fewer regulatory elements that responded in largely similar ways to immune stimuli and there is only one promoter. In contrast, there are considerably more *LAIR1* and *LAIR2* regulatory elements and they showed greater variability in response to different stimuli, suggesting potentially more complex transcriptional regulation. Consistent with this idea, the TSS/promoter element of an alternative *LAIR1* transcript had significantly higher reporter activity in myeloid cells than the canonical promoter, and RNAseq showed higher expression in CD14+ monocytes, suggesting that this alternative transcript may be expressed in myeloid cells. Notably, the alternative *LAIR1* transcript codes for a nearly identical protein sequence as the canonical transcript, suggesting that the difference in these two isoforms is related to transcriptional regulation and underscoring the conservation of the *LAIR1* amino acid sequence. In further support of this idea, Hi-C 3D genome interaction data shows that the alternative upstream promoter elements for *LAIR1* and *LAIR2* each act as a focus of interactions with several of the other regulatory elements in their respective gene loci. Together, these data point to important roles in locus organization and transcriptional regulation for the upstream promoter elements, and suggest that they may contribute to controlling the complex regulatory landscapes of *LAIR1* and *LAIR2*.

Finally, we sought to identify transcription factors (TF) that control the distinct expression patterns we detected in murine and human monocytes, BMDM, and MDM under homeostatic and stimulated conditions. Integrative analysis of TF ChIP-seq and RNA-seq revealed distinct sets of TFs with binding peak and expression profiles that correlate with LAIR gene expression and match the cell type expression patterns exhibited by LAIR genes. For example, only the *LAIR1* set includes the myeloid-predominant TF BACH1, and only the *LAIR2* set includes the T and NK cell TF, TBX21 (T-bet), which is consistent with each gene’s pattern of expression in these cell types. Because the change in LAIR expression was most robust and consistent after IFN-γ, our TF criteria included increased expression after IFN-γ treatment. As we showed, TFs identified by these criteria in all three of the LAIR gene regulatory elements are involved in inflammatory pathways that are activated by interferons, as well as by LPS, NF-κB, and cytokines (e.g., STATs, IRF2, RELA, PML, TRIM22)^43, 54–60^ or modulate response to inflammation^47, 49, 61–66^. Each TF set is distinct, yet all three contain a STAT5 protein; the *Lair1* set also contains STAT3. While STAT1 is the canonical STAT activated by IFN-γ, activation is in the form of STAT1 phosphorylation and not expression change^44^. Relevant for our findings, IFN-γ can activate also noncanonical STAT complexes as well as induce the expression of STAT3 and STAT5 via downstream signaling (e.g., via IL-6, IL-10, as well as other cytokines and growth factors)^67–71^. Though much of the literature on STAT3 and STAT5A/B has focused on their roles in oncogenesis^72–74^, these TFs also have important functions in normal, and sometimes opposing, immune functions, including cellular differentiation, cytokine response, immunity to infection, and tolerance, as evidenced by the mix of immune deficits and autoimmunity caused by germline DNA variants^58, 75–81^.

Taken together, our studies reveal epigenetic, transcriptional, and genome structural factors and processes that cooperatively or competitively regulate LAIR gene expression, resulting in patterns that are specific to species, cell-type, developmental stage, and immune stimulus. These findings provide new insights into the distinct transcriptional mechanisms by which LAIR inhibitory proteins are regulated in human and mouse myeloid cells in response to inflammatory stimuli. Moreover, they suggest that differences in LAIR gene expression between individual humans may be due to variability in TF levels and/or sequence differences in regulatory elements. Indeed, we previously demonstrated that single nucleotide variants in TF binding sites alter regulatory element activity and the expression of linked genes^82^. Such LAIR gene expression differences may impact susceptibility to immunological or infectious diseases as suggested by GWAS studies^83–86^. Future studies are needed to elucidate the gene regulatory differences that underlie distinct patterns of LAIR gene expression in different individuals and whether altered LAIR levels contribute to disease pathogenesis.

## ACKNOWLEDGEMENTS

– Conception and design: H.K. Dorando, J. Li, E. Mutic, J.E. Payton
– Development of methodology: H.K. Dorando, J. Li, E. Mutic, C. Quinn, J.E. Payton
– Acquisition of data: H.K. Dorando, J. Li, E. Mutic, E. Perrin, C. Quinn, M. Wurtz, J.E. Payton
– Analysis, interpretation of data, generation of figures: H.K. Dorando, J. Li, E. Mutic, E. Perrin, J.E. Payton
– Funding acquisition, writing, review, and/or revision of the manuscript: H.K. Dorando, J. Li, E. Mutic, J.E. Payton
– Study supervision: J.E. Payton.

This work was supported by American Heart Association grant #P23-01044 to H. Dorando, Washington University BioSURF funding to J. Li and E. Mutic, and Siteman Cancer Center Foundation and Barnes-Jewish Hospital Foundation Steinbeck Designated Fund grant support to J. Payton.

## FIGURE LEGENDS

**Figure S1.** Expression of *Lair1* across immune subsets and gating strategy for bone marrow cells and bone marrow-derived macrophages and monocytes. **A**) Expression of *Lair1* (RNAseq)^28, 29^ in immune cell subsets. **B-C)** All cells were gated based on physical parameters and a viability exclusion dye to obtain singlet, viable cells for further analysis. **(B)** Day 0; **(C)** Day 7. **D)** Number of monocytes and macrophages analyzed by flow cytometry at each time point.

**Figure S2.** Gating strategy for mouse alveolar macrophages. Plots from a representative experiment, cells from LPS-treated bronchoalveolar lavage fluid.

**Figure S3.** Monocyte-derived macrophages have low expression of *LAIR2* mRNA but expression patterns are consistent with *LAIR1* mRNA. **A-B**) LAIR1 and LAIR2 expression across immune cell subsets by single cell RNAseq^32^. **C**) Representative gating strategy shown for monocyte-derived macrophages. **D**) *LAIR2* mRNA expression as measured by qRT-PCR in MDMs after 24 hours LPS or IFNγ stimulation.

**Figure S4.** Alternative *LAIR1* and *LAIR2* transcripts with upstream promoter elements are expressed. WU Epigenome browser tracks show transcript isoforms for *LAIR1* **(A)** and *LAIR2* (**B**), homologous sequences in the mouse genome, RNAseq and ChIPseq from CD14+ monocytes and CD4+ αβ T cells^88, 89^.

**Figure S5.** Luciferase activity of every candidate enhancer element tested. Bar graphs show fold change and standard deviation of reporter activity for candidate *Lair1* enhancers (**A**), *LAIR1* enhancers (**B**), or *LAIR2* enhancers (**C**) relative to empty vector in K562 myeloid cells after 24 hours of indicated stimulations. Selected regions of interest from A-C with statistical significance are shown in Fig. 6. See **Table S5** for all statistical tests for K562 reporter assays. **D**) Fold change and standard deviation of selected elements relative to empty vector in HUT78 T cells after 24 hours of indicated stimulation. Statistical significance (two-way ANOVA with Tukey’s test for multiple comparisons): * p<0.05 for condition compared to its empty vector control; # p<0.05 for immune-stimulated condition compared to baseline for same candidate region.

**Figure S6.** 3D interactions of gene regulatory elements in *LAIR1* and *LAIR2* gene loci. UCSC tracks^87^ show candidate regulatory elements (from Figs. 5 and 7) and their 3D interactions determined by capture Hi-C genome conformation studies from the GeneHancer project^50^.

## Supplemental Tables

Supplementary Table 1. Primers for human and mouse qRT-PCR targets.

Supplementary Table 2. Flow antibodies used for human and mouse monocytes and macrophages.

Supplementary Table 3. Luciferase regions tested for mouse *Lair1* and human LAIR1&2. Genomic coordinates and forward/reverse primers listed.

Supplementary Table 4. Ct values for LAIR1 and LAIR2 qRT-PCR in human monocyte-derived macrophages.

Supplementary Table 5. Complete Luciferase two-way ANOVA statistical testing results.

## LITERATURE CITED

1. Rumpret M, Drylewicz J, Ackermans LJE, et al. Functional categories of immune inhibitory receptors. Nat Rev Immunol. Epub ahead of print July 1, 2020. DOI: 10.1038/s41577-020-0352-z.

2. Odorizzi PM, Wherry EJ. Inhibitory Receptors on Lymphocytes: Insights from Infections. J Immunol. 2012;188:2957–2965.

3. Ravetch JV, Lanier LL. Immune Inhibitory Receptors. Science. 2000;290:84–89.

4. Schnell A, Bod L, Madi A, et al. The yin and yang of co-inhibitory receptors: toward anti-tumor immunity without autoimmunity. Cell Res. 2020;30:285–299.

5. Andrews JM, Schmidt JA, Carson KR, et al. Novel cell adhesion/migration pathways are predictive markers of HDAC inhibitor resistance in cutaneous T cell lymphoma. EBioMedicine. Epub ahead of print July 26, 2019. DOI: 10.1016/j.ebiom.2019.07.053.

6. Lebbink RJ, Berg MCW van den, Ruiter T de, et al. The Soluble Leukocyte-Associated Ig-Like Receptor (LAIR)-2 Antagonizes the Collagen/LAIR-1 Inhibitory Immune Interaction. J Immunol. 2008;180:1662–1669.

7. Lebbink RJ, Ruiter T de, Adelmeijer J, et al. Collagens are functional, high affinity ligands for the inhibitory immune receptor LAIR-1. J Exp Med. 2006;203:1419–1425.

8. Meyaard L, Adema GJ, Chang C, et al. LAIR-1, a Novel Inhibitory Receptor Expressed on Human Mononuclear Leukocytes. Immunity. 1997;7:283–290.

9. Kang X, Lu Z, Cui C, et al. The ITIM-containing receptor LAIR1 is essential for acute myeloid leukaemia development. Nat Cell Biol. 2015;17:665–677.

10. Lebbink RJ, Ruiter T de, Verbrugge A, et al. The Mouse Homologue of the Leukocyte-Associated Ig-Like Receptor-1 Is an Inhibitory Receptor That Recruits Src Homology Region 2-Containing Protein Tyrosine Phosphatase (SHP)-2, but Not SHP-1. J Immunol. 2004;172:5535–5543.

11. Meyaard L, Hurenkamp J, Clevers H, et al. Leukocyte-Associated Ig-Like Receptor-1 Functions as an Inhibitory Receptor on Cytotoxic T Cells. J Immunol. 1999;162:5800– 5804.

12. Peng DH, Rodriguez BL, Diao L, et al. Collagen promotes anti-PD-1/PD-L1 resistance in cancer through LAIR1-dependent CD8 + T cell exhaustion. Nat Commun. 2020;11:4520.

13. Poggi A, Catellani S, Bruzzone A, et al. Lack of the leukocyte-associated Ig-like receptor-1 expression in high-risk chronic lymphocytic leukaemia results in the absence of a negative signal regulating kinase activation and cell division. Leukemia. 2008;22:980–988.

14. Storm L, Bruijnesteijn J, de Groot NG, et al. The Genomic Organization of the LILR Region Remained Largely Conserved Throughout Primate Evolution: Implications for Health And Disease. Front Immunol. 2021;12:716289.

15. Martin AM, Kulski JK, Witt C, et al. Leukocyte Ig-like receptor complex (LRC) in mice and men. Trends Immunol. 2002;23:81–88.

16. Keerthivasan S, Şenbabaoğlu Y, Martinez-Martin N, et al. Homeostatic functions of monocytes and interstitial lung macrophages are regulated via collagen domain-binding receptor LAIR1. Immunity. 2021;54:1511–1526.e8.

17. Jansen CA, Cruijsen CWA, de Ruiter T, et al. Regulated expression of the inhibitory receptor LAIR-1 on human peripheral T cells during T cell activation and differentiation. Eur J Immunol. 2007;37:914–924.

18. Nordkamp MJMO, Roon JAG van, Douwes M, et al. Enhanced secretion of leukocyte-associated immunoglobulin-like receptor 2 (LAIR-2) and soluble LAIR-1 in rheumatoid arthritis: LAIR-2 is a more efficient antagonist of the LAIR-1–collagen inhibitory interaction than is soluble LAIR-1. Arthritis Rheum. 2011;63:3749–3757.

19. van der Vuurst de Vries AR, Clevers H, Logtenberg T, et al. Leukocyte-associated immunoglobulin-like receptor-1 (LAIR-1) is differentially expressed during human B cell differentiation and inhibits B cell receptor-mediated signaling. Eur J Immunol. 1999;29:3160–3167.

20. Carvalheiro T, Garcia S, Pascoal Ramos MI, et al. Leukocyte Associated Immunoglobulin Like Receptor 1 Regulation and Function on Monocytes and Dendritic Cells During Inflammation. Front Immunol. 2020;11:1793.

21. Verbrugge A, Ruiter T de, Geest C, et al. Differential expression of leukocyte-associated Ig-like receptor-1 during neutrophil differentiation and activation. J Leukoc Biol. 2006;79:828–836.

22. Besteman SB, Callaghan A, Hennus MP, et al. Signal inhibitory receptor on leukocytes (SIRL)-1 and leukocyte-associated immunoglobulin-like receptor (LAIR)-1 regulate neutrophil function in infants. Clin Immunol. 2020;211:108324.

23. Chicaybam L, Barcelos C, Peixoto B, et al. An Efficient Electroporation Protocol for the Genetic Modification of Mammalian Cells. Front Bioeng Biotechnol. 2016;4:99.

24. Hammal F, de Langen P, Bergon A, et al. ReMap 2022: a database of Human, Mouse, Drosophila and Arabidopsis regulatory regions from an integrative analysis of DNA-binding sequencing experiments. Nucleic Acids Res. 2022;50:D316–D325.

25. Mostafavi S, Yoshida H, Moodley D, et al. Parsing the Interferon Transcriptional Network and Its Disease Associations. Cell. 2016;164:564–578.

26. Karlsson M, Zhang C, Méar L, et al. A single-cell type transcriptomics map of human tissues. Sci Adv. 2021;7:eabh2169.

27. The Human Protein Atlas: The blood & immune cell transcriptome Available from: https://www.proteinatlas.org/humanproteome/single+cell+type/blood+%2526+immune+cells#the_bloodimmune_cell_transcriptome.

28. ImmGen ULI: Systemwide RNA-seq profiles (#1) Available from: https://www.ncbi.nlm.nih.gov/geo/query/acc.cgi?acc=GSE109125.2020.

29. Yoshida H, Lareau CA, Ramirez RN, et al. The cis-Regulatory Atlas of the Mouse Immune System. Cell. 2019;176:897–912.e20.

30. RNAseq profiling of defined immunocyte subsets from human blood, healthy volunteers Available from: https://www.ncbi.nlm.nih.gov/geo/query/acc.cgi?acc=GSE227743. 2023.

31. Transcription profile analysis of wild type bone marrow derived-macrophages in response to Anisomycin, type I and type II interferons and the combination of them Available from: https://www.ncbi.nlm.nih.gov/geo/query/acc.cgi?acc=GSE199128. 2023.

32. Single Cell Portal: PBMCs from 2 donors Available from: https://singlecell.broadinstitute.org/single_cell/study/SCP345/ica-blood-mononuclear-cells-2-donors-2-sites?scpbr=immune-cell-atlas#study-summary.

33. Schaum N, Karkanias J, Neff NF, et al. Single-cell transcriptomics of 20 mouse organs creates a Tabula Muris. Nature. 2018;562:367–372.

34. Tabula Muris: Single-cell RNA-seq data from Smart-seq2 sequencing of FACS sorted cells (v2) Available from: https://figshare.com/articles/dataset/Single-cell_RNA-seq_data_from_Smart-seq2_sequencing_of_FACS_sorted_cells_v2_/5829687. 2018.

35. ReMap 2022 Database Available from: https://remap.univ-amu.fr/download_page.

36. Han W, Tanjore H, Liu Y, et al. Identification and Characterization of Alveolar and Recruited Lung Macrophages during Acute Lung Inflammation. J Immunol Baltim Md 1950. 2023;210:1827–1836.

37. Martínez-Esparza M, Ruiz-Alcaraz AJ, Carmona-Martínez V, et al. Expression of LAIR-1 (CD305) on Human Blood Monocytes as a Marker of Hepatic Cirrhosis Progression. J Immunol Res. 2019;2019:2974753.

38. Moore JE, Purcaro MJ, Pratt HE, et al. Expanded encyclopaedias of DNA elements in the human and mouse genomes. Nature. 2020;583:699–710.

39. Sloan CA, Chan ET, Davidson JM, et al. ENCODE data at the ENCODE portal. Nucleic Acids Res. 2016;44:D726–732.

40. Dunham I, Kundaje A, Aldred SF, et al. An integrated encyclopedia of DNA elements in the human genome. Nature. 2012;489:57–74.

41. Gerstein MB, Kundaje A, Hariharan M, et al. Architecture of the human regulatory network derived from ENCODE data. Nature. 2012;489:91–100.

42. Hoffman MM, Ernst J, Wilder SP, et al. Integrative annotation of chromatin elements from ENCODE data. Nucleic Acids Res. 2013;41:827–841.

43. Cui H, Banerjee S, Guo S, et al. IFN Regulatory Factor 2 Inhibits Expression of Glycolytic Genes and Lipopolysaccharide-Induced Proinflammatory Responses in Macrophages. J Immunol. 2018;200:3218–3230.

44. Ivashkiv LB. IFNγ: signalling, epigenetics and roles in immunity, metabolism, disease and cancer immunotherapy. Nat Rev Immunol. 2018;18:545–558.

45. Haldar M, Kohyama M, So AY-L, et al. Heme-Mediated SPI-C Induction Promotes Monocyte Differentiation into Iron-Recycling Macrophages. Cell. 2014;156:1223–1234.

46. Zhang X, Guo J, Wei X, et al. Bach1: Function, Regulation, and Involvement in Disease. Oxid Med Cell Longev. 2018;2018:1347969.

47. Kallies A, Good-Jacobson KL. Transcription Factor T-bet Orchestrates Lineage Development and Function in the Immune System. Trends Immunol. 2017;38:287–297.

48. Vicenzi E, Poli G. The interferon-stimulated gene TRIM22: A double-edged sword in HIV-1 infection. Cytokine Growth Factor Rev. 2018;40:40–47.

49. Wu J, Ma S, Hotz-Wagenblatt A, et al. Regulatory T cells sense effector T-cell activation through synchronized JunB expression. FEBS Lett. 2019;593:1020–1029.

50. Fishilevich S, Nudel R, Rappaport N, et al. GeneHancer: genome-wide integration of enhancers and target genes in GeneCards. Database J Biol Databases Curation. 2017;2017:bax028.

51. Borriello F, Iannone R, Di Somma S, et al. Lipopolysaccharide-Elicited TSLPR Expression Enriches a Functionally Discrete Subset of Human CD14+ CD1c+ Monocytes. J Immunol Baltim Md 1950. 2017;198:3426–3435.

52. Maasho K, Masilamani M, Valas R, et al. The inhibitory leukocyte-associated Ig-like receptor-1 (LAIR-1) is expressed at high levels by human naive T cells and inhibits TCR mediated activation. Mol Immunol. 2005;42:1521–1530.

53. Agashe VV, Jankowska-Gan E, Keller M, et al. Leukocyte-Associated Ig-like Receptor 1 Inhibits Th1 Responses but Is Required for Natural and Induced Monocyte-Dependent Th17 Responses. J Immunol. 2018;201:772–781.

54. Cuesta N, Salkowski CA, Thomas KE, et al. Regulation of lipopolysaccharide sensitivity by IFN regulatory factor-2. J Immunol Baltim Md 1950. 2003;170:5739–5747.

55. Kanai T, Jenks J, Nadeau KC. The STAT5b Pathway Defect and Autoimmunity. Front Immunol. 2012;3:234.

56. Khalfin-Rabinovich Y, Weinstein A, Levi B-Z. PML is a key component for the differentiation of myeloid progenitor cells to macrophages. Int Immunol. 2011;23:287–296.

57. Lukhele S, Rabbo DA, Guo M, et al. The transcription factor IRF2 drives interferon-mediated CD8+ T cell exhaustion to restrict anti-tumor immunity. Immunity. 2022;55:2369–2385.e10.

58. O’Shea JJ, Holland SM, Staudt LM. JAKs and STATs in Immunity, Immunodeficiency, and Cancer. N Engl J Med. 2013;368:161–170.

59. Reddi TS, Merkl PE, Lim S-Y, et al. Tripartite Motif 22 (TRIM22) protein restricts herpes simplex virus 1 by epigenetic silencing of viral immediate-early genes. PLoS Pathog. 2021;17:e1009281.

60. Vallabhapurapu S, Karin M. Regulation and function of NF-kappaB transcription factors in the immune system. Annu Rev Immunol. 2009;27:693–733.

61. Cain DW, Cidlowski JA. Immune regulation by glucocorticoids. Nat Rev Immunol. 2017;17:233–247.

62. Igarashi K, Kurosaki T, Roychoudhuri R. BACH transcription factors in innate and adaptive immunity. Nat Rev Immunol. 2017;17:437–450.

63. Imbratta C, Hussein H, Andris F, et al. c-MAF, a Swiss Army Knife for Tolerance in Lymphocytes. Front Immunol;11 Available from: https://www.frontiersin.org/articles/10.3389/fimmu.2020.00206. 2020. Accessed July 29, 2023.

64. Katagiri T, Kameda H, Nakano H, et al. Regulation of T cell differentiation by the AP-1 transcription factor JunB. Immunol Med. 2021;44:197–203.

65. Katsuoka F, Yamamoto M. Small Maf proteins (MafF, MafG, MafK): History, structure and function. Gene. 2016;586:197–205.

66. Yu T, Li YJ, Bian AH, et al. The Regulatory Role of Activating Transcription Factor 2 in Inflammation. Mediators Inflamm. 2014;2014:e950472.

67. Barrat FJ, Crow MK, Ivashkiv LB. Interferon target-gene expression and epigenomic signatures in health and disease. Nat Immunol. 2019;20:1574–1583.

68. Platanias LC. Mechanisms of type-I– and type-II-interferon-mediated signalling. Nat Rev Immunol. 2005;5:375–386.

69. Sarma U, Maiti M, Nair A, et al. Regulation of STAT3 signaling in IFNγ and IL10 pathways and in their cross-talk. Cytokine. 2021;148:155665.

70. Suarez AAR, Renne NV, Baumert TF, et al. Viral manipulation of STAT3: Evade, exploit, and injure. PLOS Pathog. 2018;14:e1006839.

71. Villarino AV, Kanno Y, O’Shea JJ. Mechanisms and consequences of Jak–STAT signaling in the immune system. Nat Immunol. 2017;18:374–384.

72. Johnson DE, O’Keefe RA, Grandis JR. Targeting the IL-6/JAK/STAT3 signalling axis in cancer. Nat Rev Clin Oncol. 2018;15:234–248.

73. Waldmann TA, Chen J. Disorders of the JAK/STAT Pathway in T Cell Lymphoma Pathogenesis: Implications for Immunotherapy. Annu Rev Immunol. 2017;35:533–550.

74. Wingelhofer B, Neubauer HA, Valent P, et al. Implications of STAT3 and STAT5 signaling on gene regulation and chromatin remodeling in hematopoietic cancer. Leukemia. 2018;32:1713–1726.

75. Casanova J-L, Holland SM, Notarangelo LD. Inborn errors of human JAKs and STATs. Immunity. 2012;36:515–528.

76. Haapaniemi EM, Kaustio M, Rajala HLM, et al. Autoimmunity, hypogammaglobulinemia, lymphoproliferation, and mycobacterial disease in patients with activating mutations in STAT3. Blood. 2015;125:639–648.

77. Kollmann S, Grausenburger R, Klampfl T, et al. A STAT5B–CD9 axis determines self-renewal in hematopoietic and leukemic stem cells. Blood. 2021;138:2347–2359.

78. Korenfeld D, Roussak K, Dinkel S, et al. STAT3 Gain-of-Function Mutations Underlie Deficiency in Human Nonclassical CD16+ Monocytes and CD141+ Dendritic Cells. J Immunol. 2021;207:2423–2432.

79. Milner JD, Vogel TP, Forbes L, et al. Early-onset lymphoproliferation and autoimmunity caused by germline STAT3 gain-of-function mutations. Blood. 2015;125:591–599.

80. Stark GR, Darnell JE. The JAK-STAT Pathway at Twenty. Immunity. 2012;36:503–514.

81. Villarino A, Laurence A, Robinson GW, et al. Signal transducer and activator of transcription 5 (STAT5) paralog dose governs T cell effector and regulatory functions. eLife. 2016;5:e08384.

82. Koues OI, Kowalewski RA, Chang L-WW, et al. Enhancer Sequence Variants and Transcription-Factor Deregulation Synergize to Construct Pathogenic Regulatory Circuits in B-Cell Lymphoma. Immunity. 2015;42:186–198.

83. Achieng AO, Hengartner NW, Raballah E, et al. Integrated OMICS platforms identify LAIR1 genetic variants as novel predictors of cross-sectional and longitudinal susceptibility to severe malaria and all-cause mortality in Kenyan children. EBioMedicine. 2019;45:290– 302.

84. Achieng AO, Guyah B, Cheng Q, et al. Molecular basis of reduced LAIR1 expression in childhood severe malarial anaemia: Implications for leukocyte inhibitory signalling. EBioMedicine. 2019;45:278–289.

85. Camargo CM, Augusto DG, Petzl-Erler ML. Differential gene expression levels might explain association of LAIR2 polymorphisms with pemphigus. Hum Genet. 2016;135:233– 244.

86. Farias TDJ, Augusto DG, de Almeida RC, et al. Screening the full leucocyte receptor complex genomic region revealed associations with pemphigus that might be explained by gene regulation. Immunology. 2019;156:86–93.

87. Nassar LR, Barber GP, Benet-Pagès A, et al. The UCSC Genome Browser database: 2023 update. Nucleic Acids Res. 2023;51:D1188–D1195.

88. Chadwick LH. The NIH Roadmap Epigenomics Program data resource. Epigenomics. 2012;4:317–24.

89. Li D, Purushotham D, Harrison JK, et al. WashU Epigenome Browser update 2022. Nucleic Acids Res. 2022;50:W774–W781.

