## Supplementary figures and images for "Epigenetic regulation of Leukocyte associated immunoglobulin-like receptors 1 and 2 by interferon signaling in macrophages and T cells"

### Supplemental_figures1-6

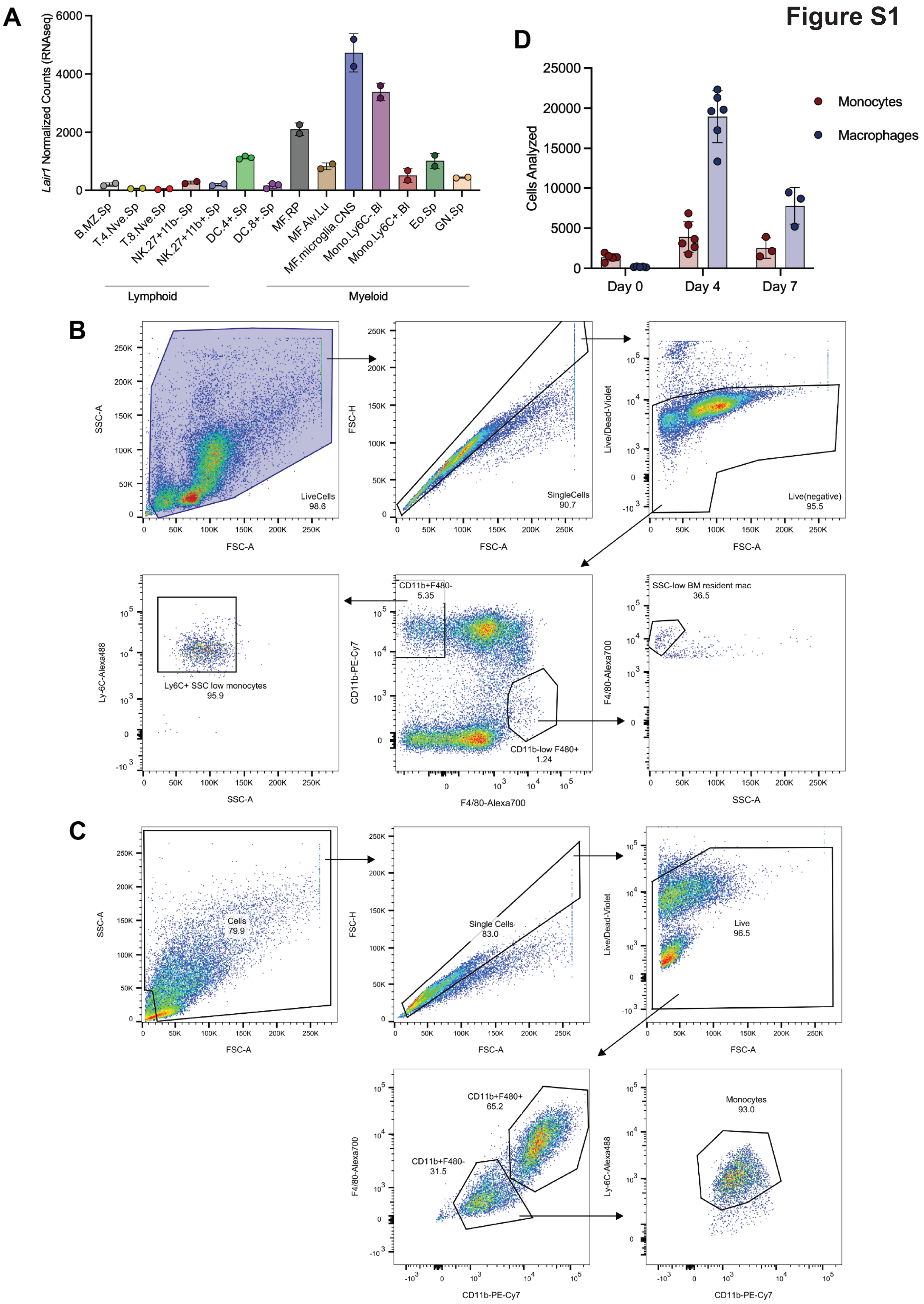

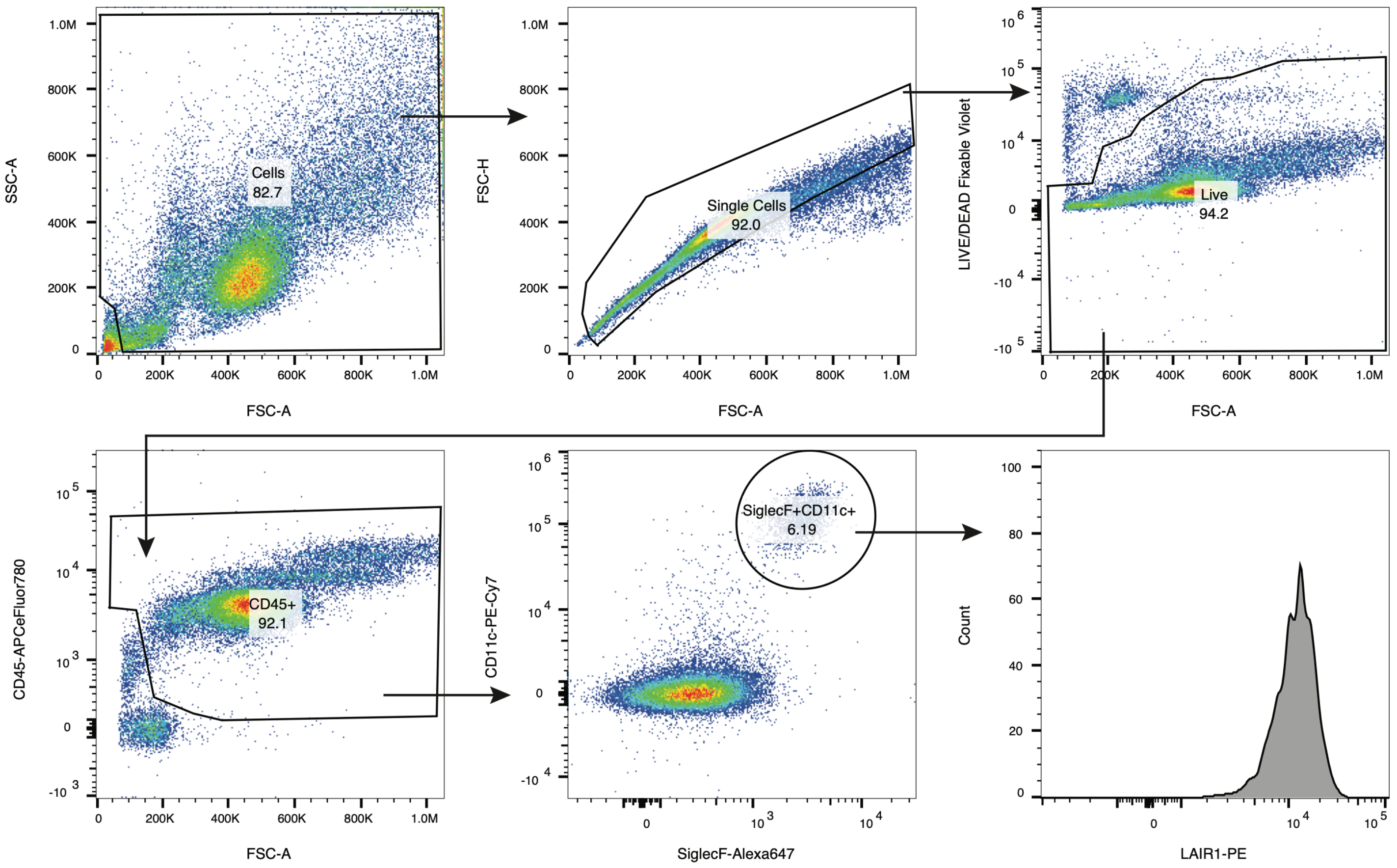

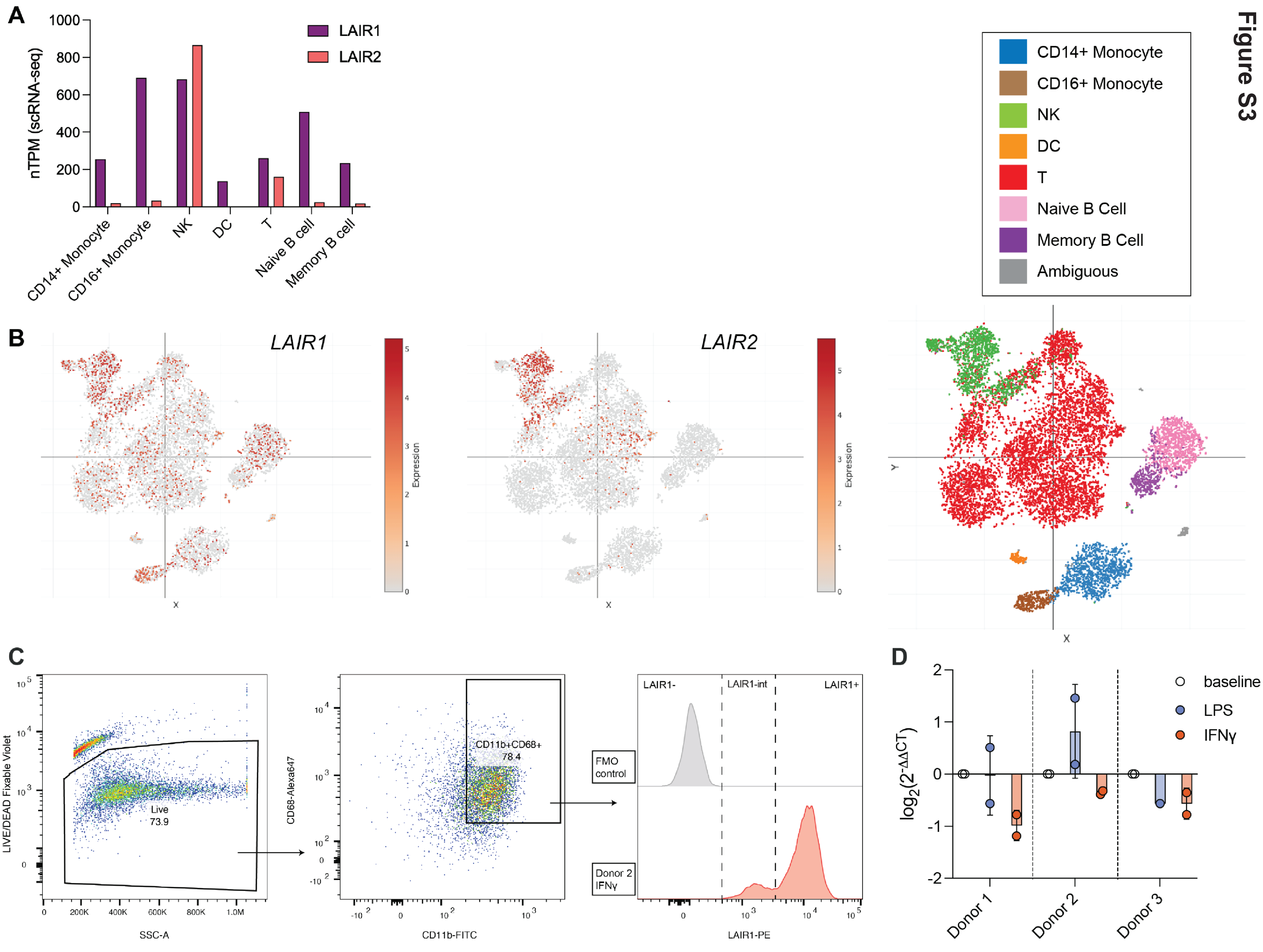

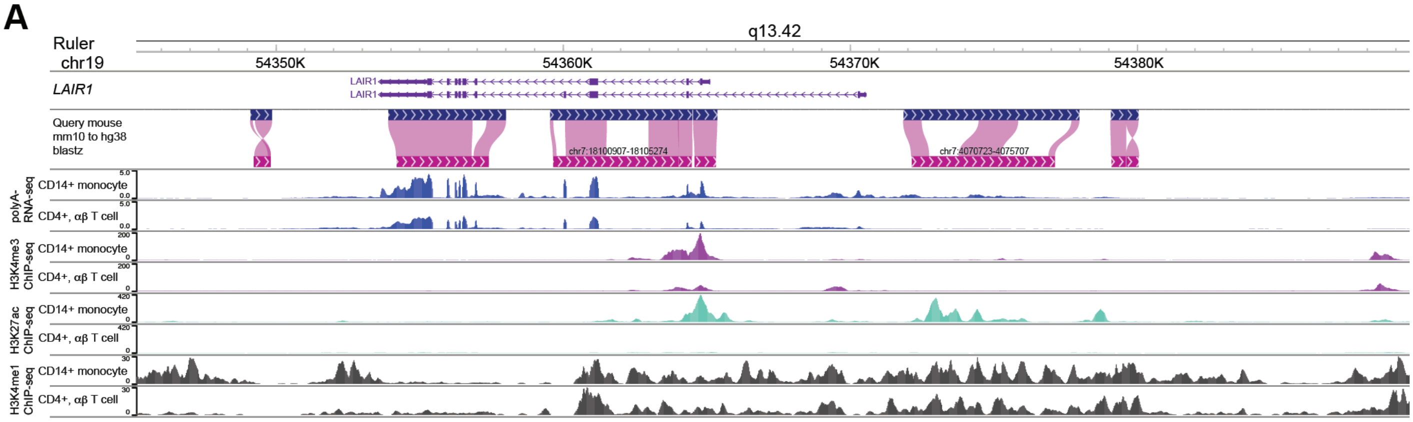

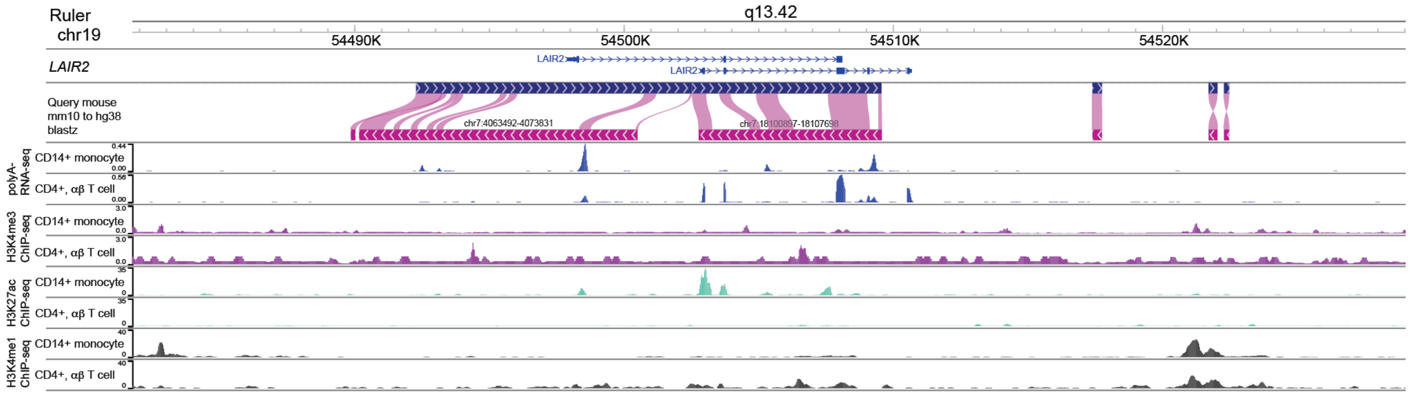

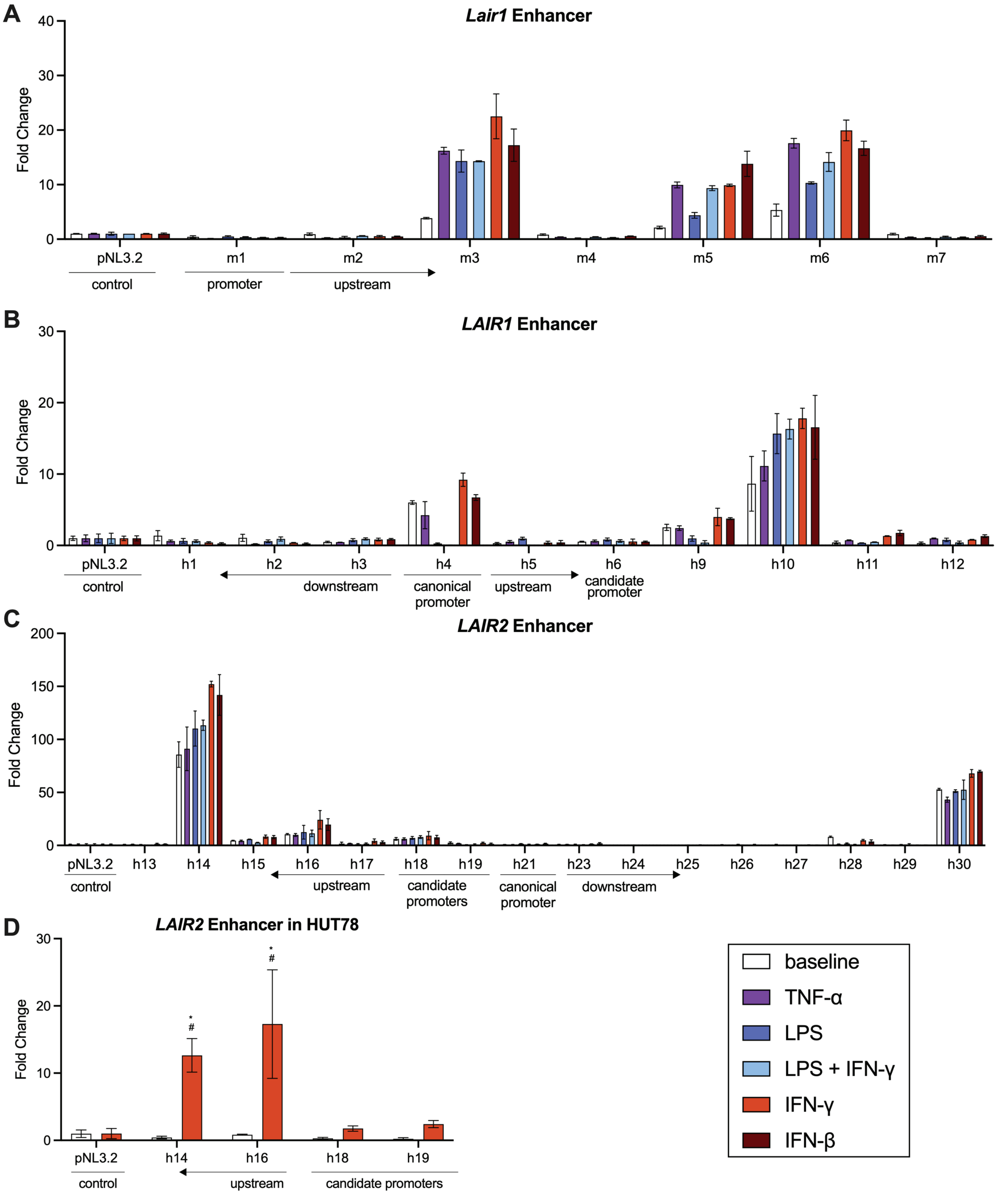

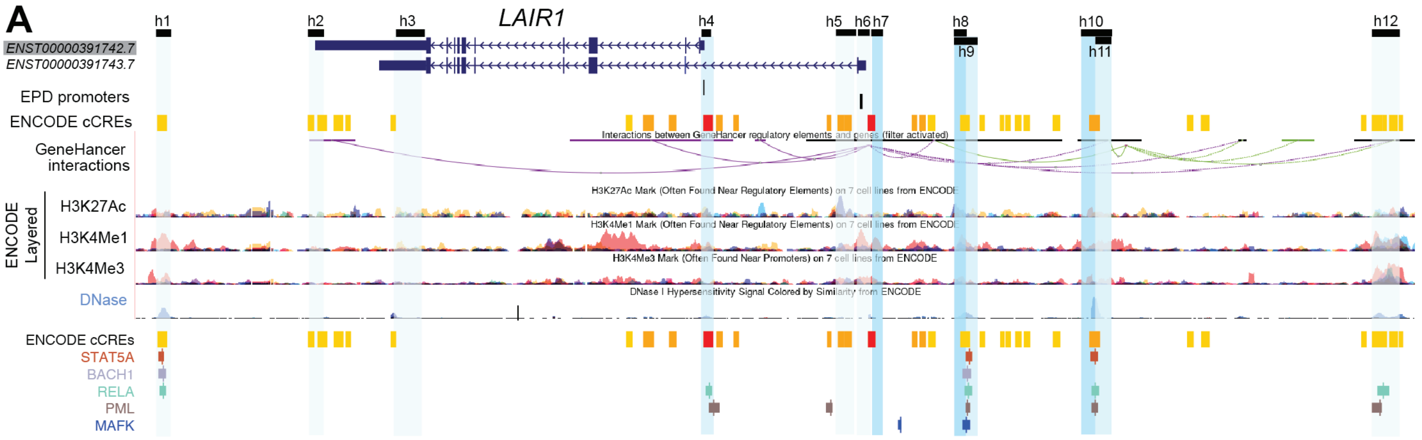

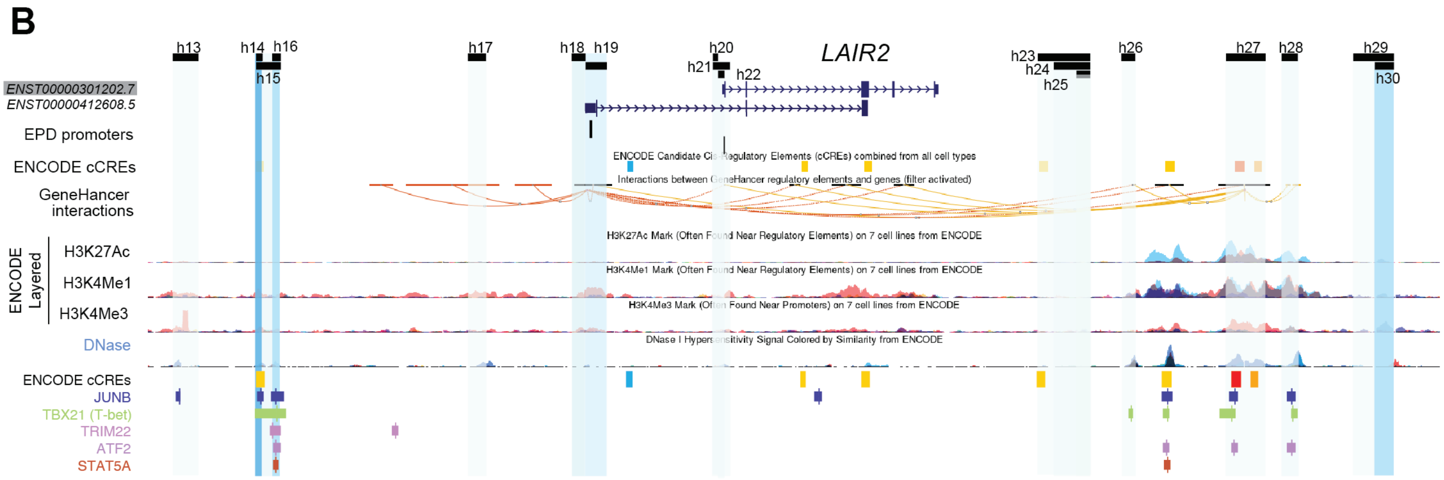
