## Supplementary material for "Epigenetic regulation of Leukocyte associated immunoglobulin-like receptors 1 and 2 by interferon signaling in macrophages and T cells": Tables1-4

**Supplementary Table 1.** Primers for human and mouse qRT-PCR targets.

| Target | Species | F Primer (5' --> 3') | R Primer (5' --> 3') |
|---|---|---|---|
| *Lair1* | mouse | ATTAACACACAGGAGGGTTCTC | CTCCAGGCGGACCATGTTA |
| *Actb* | mouse | GATGTATGAAGGCTTTGGTC | TGTGCACTTTTATTGGTCTC |
| *ACTB* | human | CTGGCACCCAGCACAATGA | AAGTCATAGTCCGCCTAGAAGC |
| *LAIR1* | human | GTCGGACAACAGTCACAATGAG | TGCTCCTCGTCCTTGCTTCT |
| *LAIR2* | human | TGGGCCTAGTGCTCTGC | TCTATCCTCCCTCTCCAGGC |

**Supplementary Table 2.** Flow antibodies used for human and mouse monocytes and macrophages.

| Species | Marker | Clone | Conjugate | Supplier | Catalog # |
|---|---|---|---|---|---|
| mouse | anti-CD11b | M1/70 | PE-Cyanine7 | BioLegend | 101215 |
| mouse | anti-F4/80 | BM8 | Alexa Fluor 700 | BioLegend | 123129 |
| mouse | anti-Ly6C | HK1.4 | Alexa Fluor 488 | Invitrogen | 53-5932-82 |
| mouse | anti-LAIR1 | 113 | PE | Invitrogen | 12-3051-82 |
| mouse | anti-CD45 | 30-F11 | APC-eFluor780 | Invitrogen | 501129642 |
| mouse | anti-Siglec-F | AB_2687570 | Alexa Fluor 647 | BD Biosciences | 562680 |
| mouse | anti-CD11c | N418 | PE-Cy7 | Invitrogen | 25-0114-82 |
| human | anti-CD11b | ICRF44 | FITC | Invitrogen | 11-0118042 |
| human | anti- CD68 | Y1/82A | Alexa Fluor 647 | Invitrogen | 51-0689-42 |
| human | anti-LAIR1 | NKTA255 | PE | Invitrogen | 12-3059-42 |

**Supplementary Table 3.** Luciferase regions tested for *Lair1*, *LAIR1*, and *LAIR2*.

| ***Lair1*** | | | | | |
|---|---|---|---|---|---|
| Chr | Start | End | Name | F Primer (5' --> 3') | R Primer (5' --> 3') |
| chr7 | 4062994 | 4063336 | m1 | CCTTCGCTGCTCTCAAGGC | TCCATGTGGGGAACCTTAGC |
| chr7 | 4073232 | 4073645 | m2 | GCACCTGATTTTGCAGCCTC | AGGCATGTCAAGCACAGTGATA |
| chr7 | 4080366 | 4081058 | m3 | GGAGTTGGGCTGGTTTCTCA | ACCACAGGATGGCTTGTCAG |
| chr7 | 4080370 | 4080760 | m4 | AAACGGAGTTGGGCTGGTTT | AAACATGCCAAGCCAACGTG |
| chr7 | 4082974 | 4083140 | m5 | CTGGGGTTTGCCCATTCTGA | GACCTCATCTCCTGCTGCAT |
| chr7 | 4082973 | 4083156 | m6 | TCTGGGGTTTGCCCATTCTG | AGTCCACCTAGCTGAGGACC |
| chr7 | 4084202 | 4084412 | m7 | TGCTACCTCTAACCTGCCCT | ACAGACGTGAAGTTGGCACA |
| ***LAIR1*** | | | | | |
| Chr | Start | End | Name | F Primer (5' --> 3') | R Primer (5' --> 3') |
| chr19 | 54345825 | 54346360 | h1 | AGTCAACTTTCACGCAGACC | ACTTCTGTGCCAGTGACTTCC |
| chr19 | 54351124 | 54351697 | h2 | GAAAATTCCCATCCCGCAGC | AGGAGCAAGAGCGTGTTTCA |
| chr19 | 54354072 | 54355068 | h3 | TGAGATTCTGCCGCCTCCTA | AATCGAGCAGCTCCTTGGAC |
| chr19 | 54364835 | 54365179 | h4 | GAGGCGCACCAATGCAAGG | AAGAGCTGCGACCGTTAACTT |
| chr19 | 54369318 | 54370116 | h5 | TTCCCCTAGTATCTGAACTCCA | CAAGAAGTACAGACCTCCGGG |
| chr19 | 54370276 | 54370696 | h6 | ATCTTCTGTCGCGGATGCAA | TCGGAGACGTATACAGGGCA |
| chr19 | 54370762 | 54371172 | h7 | TGCTTGAGAGCCAAGGCAAT | AGGTAGGAGGGGGAAACAGG |
| chr19 | 54373586 | 54374042 | h8 | TGGGAACAGGGTCAAATAGCAA | TACCCCCACCAAGGGACTTT |
| chr19 | 54373586 | 54374414 | h9 | TGGGAACAGGGTCAAATAGCAA | CTAAGGGAACTGGGAGGGGA |
| chr19 | 54377890 | 54378943 | h10 | TGATGGAAGTTTGGCTTCCAGA | TCGTGTCTGCCTTTCTCTTTCT |
| chr19 | 54378393 | 54378943 | h11 | AGGAAACCAAGCAAGGGGAC | TCGTGTCTGCCTTTCTCTTTCT |
| chr19 | 54387993 | 54388967 | h12 | CGACGCTCAACTTTTCACCC | AGCGAGAGCTTCATAGCCAC |
| ***LAIR2*** | | | | | |
| Chr | Start | End | Name | F Primer (5' --> 3') | R Primer (5' --> 3') |
| chr19 | 54482894 | 54483834 | h13 | AGAGCCAGACTCCATCCCAA | GCCCCCTTTAAGGAGCTGAG |
| chr19 | 54485916 | 54486151 | h14 | ACTGACTGATGAGAAGTCTGATGT | TGCAACTTGAGGAGCTGGAA |
| chr19 | 54485916 | 54486809 | h15 | TGACCACTTCTTAATCCACCAAG | TGACCACTTCTTAATCCACCAAG |
| chr19 | 54486524 | 54486809 | h16 | GTGTGCTTAAAACAATCCACATTTC | TGACCACTTCTTAATCCACCAAG |
| chr19 | 54493631 | 54494278 | h17 | AGTGGGAGTCCAGGAAAACG | TGGTCAGTCACACATGGATGC |
| chr19 | 54497410 | 54497885 | h18 | GCCTCTGATTTCCCTGGAGC | TAAGAAGCTCCAACCGCAGG |
| chr19 | 54497918 | 54498680 | h19 | ACCACATCCTGTGCGGTTTT | ATTTCAAAACTATTCCAGCACCATC |
| chr19 | 54502520 | 54502719 | h20 | TGGGGAAGGATTTCTATGACTCT | TGACCTTAAACTAGTTACCCGATGT |
| chr19 | 54502520 | 54503165 | h21 | TGGGGAAGGATTTCTATGACTCT | AAAGCTGACCTCATCCCCAC |
| chr19 | 54502721 | 54502937 | h22 | CTGATCATGCGGGTAAGCGA | CAGTGAGGTGTGGAGACATGG |
| chr19 | 54514331 | 54516264 | h23 | AGCATCCCTCCTTCTGTTGC | AATCTCCAGTCACTGCCAGC |
| chr19 | 54514933 | 54516264 | h24 | TTTCCTTGGGGATTCGGGTG | AATCTCCAGTCACTGCCAGC |
| chr19 | 54515767 | 54516264 | h25 | ATGGAGGGATCACAGACCCA | AATCTCCAGTCACTGCCAGC |
| chr19 | 54517394 | 54517890 | h26 | AGGCAAGGCCAGAGTGATTC | CGCCTTAACAACTGGCATCC |
| chr19 | 54521178 | 54522617 | h27 | TTTGTTCACGGTGTGGGGAG | GAGCCCTGCTAGTCTAGGGGAT |
| chr19 | 54523211 | 54523816 | h28 | ACGCCAGGTATTCCAGATGC | CATGGCCTCCTTGACACCTT |
| chr19 | 54525804 | 54527292 | h29 | CTGAAGCCTCTGATGCCGTT | GGAGCCATCGCATGGAAGAT |
| chr19 | 54526588 | 54527292 | h30 | CAGGTCTCAGCATGGTCGAA | GGAGCCATCGCATGGAAGAT |

**Supplementary Table 4**. $C_T$ values for LAIR1 and LAIR2 qRT-PCR in human monocyte-derived macrophages.

| **mean $C_T$** | **Donor 1** | | **Donor 2** | | **Donor 3** | |
|---|---|---|---|---|---|---|
| | replicate 1 avg | replicate 2 avg | replicate 1 avg | replicate 2 avg | replicate 1 avg | replicate 2 avg |
| **ACTB** | 19.48 | 15.86 | 15.20 | 16.52 | 16.97 | 15.78 |
| **LAIR1** | 25.17 | 19.65 | 20.63 | 20.19 | 21.60 | 18.86 |
| **LAIR2** | 29.17 | 22.08 | 24.26 | 22.02 | 25.42 | 21.26 |
